# Role of Macrophage PARP1 in the Regulation of Crosstalk between Adipose Immune Cells and Adipocytes during Diet-induced Obesity

**DOI:** 10.1101/2025.06.20.660762

**Authors:** Jingbo Zhu, Rebecca Gupte, Tulip Nandu, Kai Huang, W. Lee Kraus, Dan Huang

**Affiliations:** Department of Cardiology, Union Hospital, Tongji Medical College, Huazhong University of Science and Technology, Wuhan, Hubei Province 430022, P. R. China; Clinic Center of Human Gene Research, Union Hospital, Tongji Medical College, Huazhong University of Science and Technology, Wuhan, Hubei Province 430022, P. R. China; Cecil H. and Ida Green Center for Reproductive Biology Sciences, University of Texas Southwestern Medical Center, Dallas, TX 75390, USA; Computational Core Facility, Cecil H. and Ida Green Center for Reproductive Biology Sciences, University of Texas Southwestern Medical Center, Dallas, TX 75390, USA; Section of Laboratory Research, Department of Obstetrics and Gynecology, University of Texas Southwestern Medical Center, Dallas, TX 75390, USA

**Author notes:** Current address: Freenome, South San Francisco, CA 94005. Corresponding authors: Dan Huang and W. Lee Kraus. Lead Contact (for manuscript correspondence): Dan Huang, Ph.D., M.D., Cecil H. and Ida Green Center for Reproductive Biology Sciences The University of Texas Southwestern Medical Center at Dallas 5323 Harry Hines Boulevard, MC8511, Dallas, TX 75390-8511. **<u>Disclosures</u>**: W.L.K. is a holder of U.S. patent number 9,599,606, covering the ADP-ribose detection reagents used here, which have been licensed to and are sold by MilliporeSigma.

**Keywords:** PARP1, Macrophage, Adipose Tissue, Adipogenesis, Metabolism, Inflammation, NK cell, Obesity, ADP-ribosylation, Post-translational modification

## Abstract

Adipose tissue consists of heterogeneous cell populations, including macrophages, which play a key role in maintaining adipose tissue homeostasis. We previously identified PARP1 as a critical regulator of proadipogenic gene expression in preadipocytes and proinflammatory gene expression in macrophages. To investigate the role of macrophage PARP1 in regulating adipose tissue homeostasis, we generated myeloid lineage-specific *Parp1* knockout mice (*Parp1* KO^LysM^). When subjected to a high fat diet for 12 weeks, the *Parp1* KO^LysM^ mice exhibited an obese phenotype accompanied by white adipose tissue (WAT) dysfunction, characterized by altered metabolite profile, pronounced adipocyte hypertrophy, and increased macrophage infiltration. Coculture of primary preadipocytes with the conditioned medium from bone marrow-derived macrophages (BMDMs) isolated from *Parp1* KO^LysM^ or control mice demonstrated that macrophage PARP1 depletion inhibited LPS-induced proinflammatory gene expression in BMDMs, but enhanced differentiation of preadipocytes into mature adipocytes. Single cell RNA-sequencing using CD45^+^-sorted WAT resident immune cells showed that macrophage PARP1 depletion increased the fraction of macrophages and NK cells, altered gene expression in both cell populations, and promoted intercellular communications. Taken together, our studies demonstrate a key role for macrophage PARP1 in the maintenance of adipose tissue inflammatory and metabolic homeostasis. Macrophage PARP1 depletion promotes cell-cell crosstalk among macrophages, fat cells, and other immune cell populations in adipose tissue, which cooperatively drives the development of obesity.

**Significance:** In this study, we characterized a role for macrophage PARP1 in regulating adipose tissue homeostasis. Depletion of PARP1 in macrophages enhances cell-cell crosstalk among immune cells and fat cells, which exacerbates adipose dysfunction and drives obesity and adverse metabolic outcomes.

## Introduction

Obesity is a global public health issue, contributing significantly to the development of metabolic complications, such as type II diabetes, cardiovascular disease, and fatty liver disease [1]. White adipose tissue (WAT), the primary fat-storing reservoir, is crucial for maintaining whole-body energy homeostasis. The impaired function of WAT is strongly associated with obesity and its related metabolic consequences. Maintenance of WAT homeostasis involves heterogeneous cell populations in WAT, including adipocytes, adipocyte precursors, immune cells, stem cells, fibroblasts, endothelial cells, and vascular smooth muscle cells [2]. Cell-cell crosstalk may initiate and propagate WAT dysfunction. The mechanisms linking WAT dysfunction with obesity involve distinct cell types, such as altered cellular composition, adipocyte hypertrophy, preadipocyte differentiation, and the immune responses driven by various immune cell populations, among which adipose tissue macrophages play a significant role in promoting chronic low-grade inflammation during the development of obesity and metabolic dysfunction [3, 4]. However, previous studies typically focused on a single cellular component, without exploring the inter-relationships between different cell types. In addition, the regulatory mechanisms underlying cell-cell crosstalk in adipose tissue during the development of obesity is still less understood.

Poly(ADP-ribose) polymerase (PARP) 1, the founding member of the PARP family, catalyzes poly(ADP-ribosyl)ation (PARylation), an NAD^+^-dependent post-translational modification of proteins [5–7]. Although initial studies were focused on the biochemistry and molecular biology of PARP1 in DNA repair [8], new findings have demonstrated diverse roles for PARP1 in biological processes, such as metabolism, inflammation, immunity, and stress responses [5, 9–11]. Emerging evidence, including work from our group, has demonstrated that PARP1 plays key roles in metabolic physiology and disorders, including adipogenesis, obesity, hyperlipidemia, type II diabetes, and fatty liver diseases [11–21]. However, the results from these studies have not always been consistent, with some reporting contradictory results. In some cases, this may be due to the use of whole-body *Parp1* knockout or pharmacological inhibition of PARP1 activity in animal models, since PARP1 may play opposing roles in different metabolic tissues [22]. By using a preadipocyte-specific *Parp1* knockout and lineage tracing mouse genetic model, we recently showed that PARP1 acts in adipocyte precursors to control adipogenesis through PARylation of histone H2B, preventing diet-induced obesity [11]. We also showed that PARP1 acts as a key regulator of inflammatory responses in macrophages, through PARylation of the transcription factor STAT1α [10]. However, crosstalk between macrophages and adipocytes, and the contribution of macrophage-dependent PARP1 in adipogenesis, has not been characterized.

Herein, we generated a macrophage-specific *Parp1* knockout genetic mouse model. Using this model, we found that macrophage PARP1 is required to maintain optimal inflammatory response and metabolic homeostasis of WAT. Together, our data demonstrate that deletion of PARP1 in macrophage exacerbates WAT dysfunction through enhanced adipogenesis and immune cell crosstalk during the development of obesity.

## Results

### PARP1 depletion in macrophages promotes metabolic alterations and the development of obesity

To investigate the role of macrophage PARP1 in the control of body weight and metabolic homeostasis, we deleted the *Parp1* gene in the myeloid lineage by crossing *Parp1^loxp/loxp^* mice with *LysM-Cre* mice to generate myeloid lineage-specific *Parp1* knockout mice (i.e., *Parp1 ^loxp/loxp^*/*LysM-Cre^tg/-^*, hereafter referred to as *Parp1* KO^LysM^) (Figure 1A). *Parp1^loxp/loxp^*/*LysM-Cre^-/-^* littermates were used for comparison as a control. To investigate the function of macrophage PARP1 in regulating metabolic outcomes, we fed eight-week old male *Parp1* KO^LysM^ and control mice with a high fat diet (HFD; 60% of the total calories derived from fat) (Figure 1B). After 12-week HFD feeding, *Parp1* KO^LysM^ mice exhibited a significant increase in body weight compared to control mice (Figure 1, C and D). In addition, we observed a significantly higher fat mass in *Parp1* KO^LysM^ mice than control mice by using an EchoMRI-100 body composition analyzer (Figure 1E).

**Figure 1.**
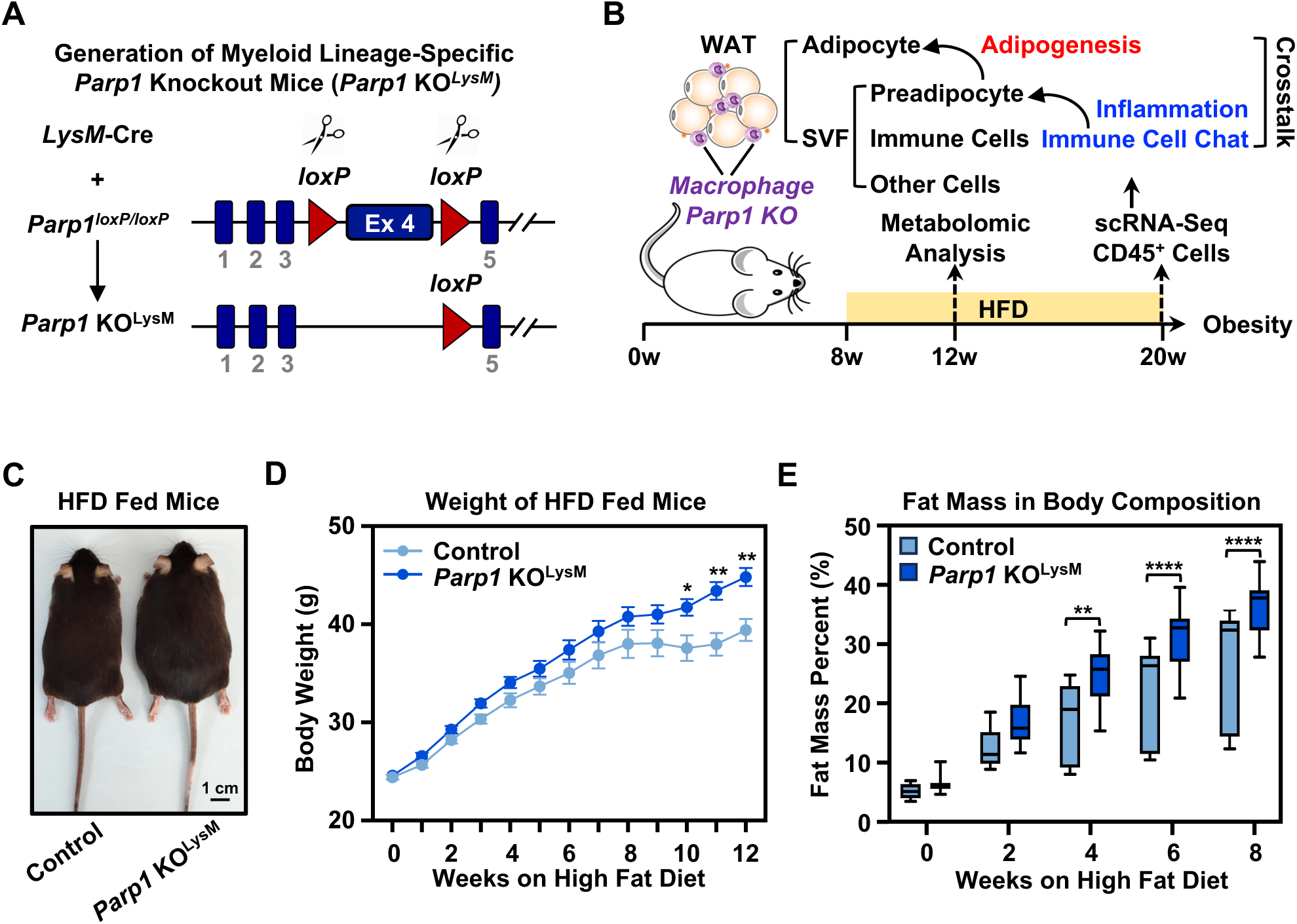
Macrophage-specific depletion of PARP1 promotes diet-induced obesity. **(A)** Schematic representation of generation of myeloid lineage-specific *Parp1* knockout mice (*Parp1* KO^LysM^) by crossing *Parp1^loxp/loxp^* mice with *LysM-Cre* mice. **(B)** Schematic representation of the design of HFD-induced obesity in mice. Eight-week old control and *Parp1* KO^LysM^ mice were fed with HFD. After 4 weeks of HFD feeding, WAT metabolomics was analyzed. After 12 weeks of HFD feeding, scRNA-seq was performed using sorted CD45^+^ immune cells from SVF of WAT. WAT was collected and assayed. **(C)** Representative images of control and *Parp1* KO^LysM^ mice after 12 weeks of HFD feeding. **(D)** Body weights of control and *Parp1* KO^LysM^ mice measured weekly following the start of HFD. Each point represents the mean ± SEM (n = 18 for control group, and n = 23 for *Parp1* KO^LysM^ group). Asterisks indicate significant differences from the control at individual time points; two tailed, unpaired t-test; *, p < 0.05; and **, p < 0.01. **(E)** Box plots showing fat mass (normalized to body weight) of control and *Parp1* KO^LysM^ mice measured every two weeks following the start of HFD (n = 8 for each group). Asterisks indicate significant differences from the control at individual time points; two tailed, unpaired t-test; **, p < 0.01; and ****, p < 0.0001.

To investigate the effects of macrophage PARP1 depletion on energy homeostasis, we analyzed energy expenditure of control and *Parp1* KO^LysM^ mice on a HFD by using a comprehensive laboratory animal monitoring system (CLAMS). In comparison to control mice, *Parp1* KO^LysM^ mice exhibited lower energy expenditure as demonstrated by significant decreases in oxygen consumption (Figure 2A), carbon dioxide production (Figure 2B), and heat generation (Figure 2C). These results indicate that macrophage-specific deletion of PARP1 exacerbates diet-induced obesity in mice. As obesity is strongly linked to metabolic dysfunction in the liver, we then examined livers from HFD-fed control and *Parp1* KO^LysM^ mice. Compared to control mice, *Parp1* KO^LysM^ mice exhibited a significant increase in liver weight (Figure 2D), accompanied by an increase in the expression of hepatic de novo lipogenesis genes, including *Srebp1, Scd1,* and *Fasn* (Figure 2E), which are related to the development of non-alcoholic fatty liver disease (NAFLD). Consistence with these gene expression results, hematoxylin and eosin (H&E) staining of the liver sections demonstrated enhanced lipid accumulation in *Parp1* KO^LysM^ mice versus control mice (Supplementary Figure 1A). However, glucose tolerance tests (GTT) (Supplementary Figure 1B), insulin tolerance tests (ITT) (Supplementary Figure 1C), and serum lipid analyses (Supplementary Figure 1D) showed no significant differences between control and *Parp1* KO^LysM^ mice. These results suggest that depletion of PARP1 in macrophage promotes the development of obesity and related metabolic consequences, such as NAFLD.

**Figure 2.**
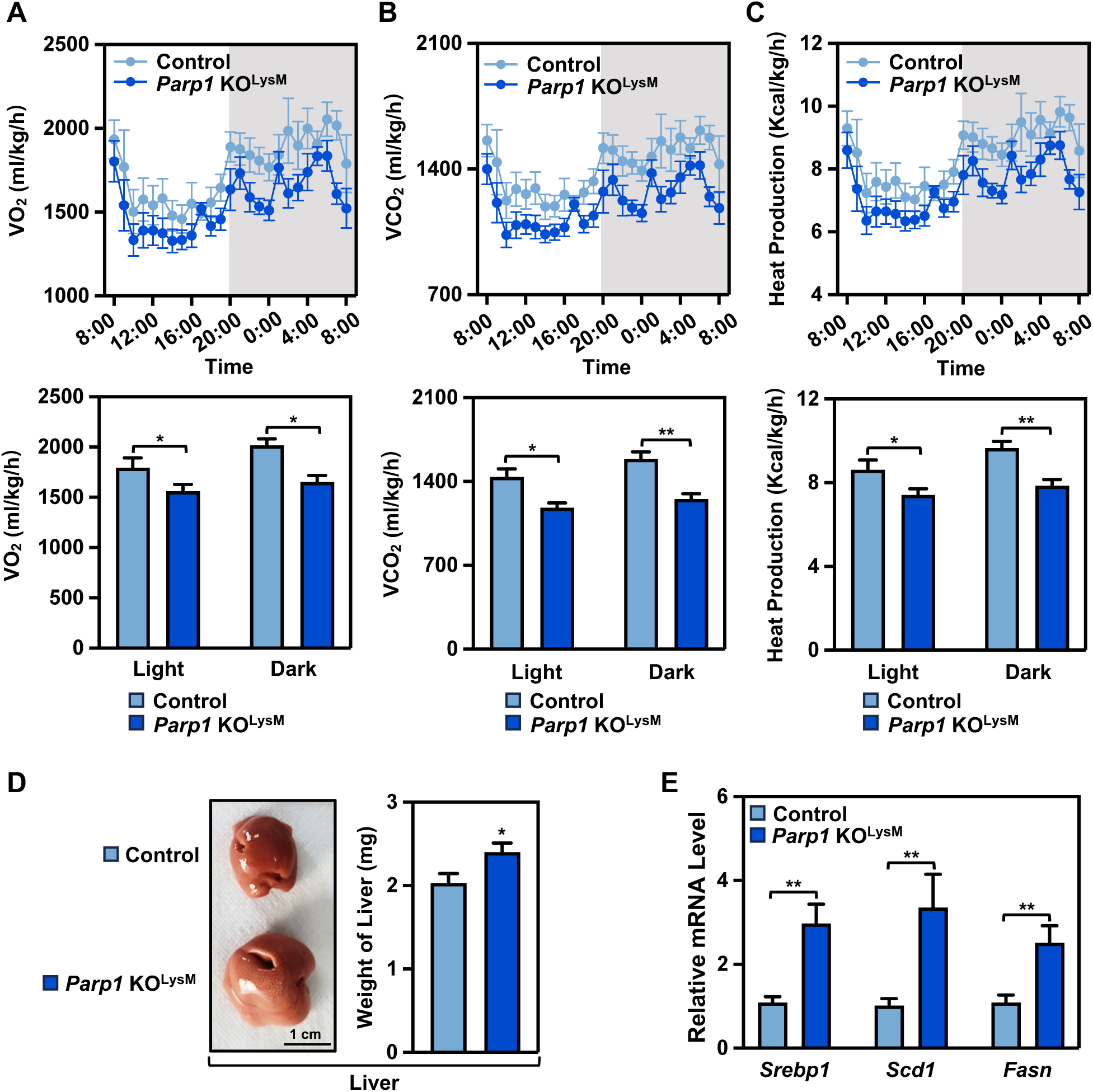
PARP1 depletion in macrophages promotes obesity-associated metabolic alterations. **(A-C)** Analysis (*Upper panels*) and quantification (*lower panels*) of oxygen consumption (**A**), carbon dioxide production (**B**), and heat generation **(C)** in control and *Parp1* KO^LysM^ mice after 12 weeks of HFD feeding, by using a comprehensive laboratory animal monitoring system. Each point in the line graphs represents the mean ± SEM. Each bar represents the mean + SEM during a 12 h-12 h light-dark cycle. (n = 8 for each group). Asterisks indicate significant differences from the control; two tailed, unpaired t-test; *, p < 0.05; and **, p < 0.01. **(D)** Representative images of liver (*Left*) and weight of liver (*Right*) from control and *Parp1* KO^LysM^ mice after 12 weeks of HFD feeding. Each bar represents the mean + SEM (n = 15 for each group). Asterisks indicate significant differences from the control; two tailed, unpaired t-test; *, p < 0.05. **(E)** Expression of de novo lipogenesis genes in liver, including *Srebp1, Scd1,* and *Fasn,* as determined by RT-qPCR. Each bar represents the mean + SEM (n = 10 for each group). Asterisks indicate significant differences from the control; two tailed, unpaired t-test; **, p < 0.01.

### Depletion of PARP1 in macrophages alters metabolite profile of white adipose tissue

As white adipose tissue (WAT) is a critical energy reservoir for maintaining systemic energy homeostasis, diet-induced imbalances between energy intake and expenditure increases adiposity and leads to obesity through accumulation of WAT [23, 24]. To determine the changes in the metabolic profile of WAT in the development of obesity, we performed untargeted metabolomics for WAT from control or *Parp1* KO^LysM^ mice after 4 weeks of HFD feeding, when *Parp1* KO^LysM^ mice started showing greater weight gain. A total of 113 differentially accumulated metabolites (DAMs) were identified in *Parp1* KO^LysM^ mice versus control mice, of which 67 were upregulated and 46 were downregulated (Figure 3, A and B). KEGG pathway analysis demonstrated that the DAMs were mainly enriched in metabolic pathways, involved in glycerophospholipid metabolism, polyunsaturated fatty acid metabolism (including arachidonic acid, alpha-linolenic acid, and linoleic acid), and glycerolipid biosynthesis pathways (Figure 3C). The DAMs were classified into different categories, of which the most enriched category was glycerophospholipid (Figure 3D). Overall, the metabolomics analysis indicates that depletion of PARP1 in macrophages alters the metabolite profile in WAT, mainly through the remodeling of glycerophospholipid metabolism and polyunsaturated fatty acid metabolism, which may affect adipocyte function, inflammation state, and energy balance of WAT [25–27].

**Figure 3.**
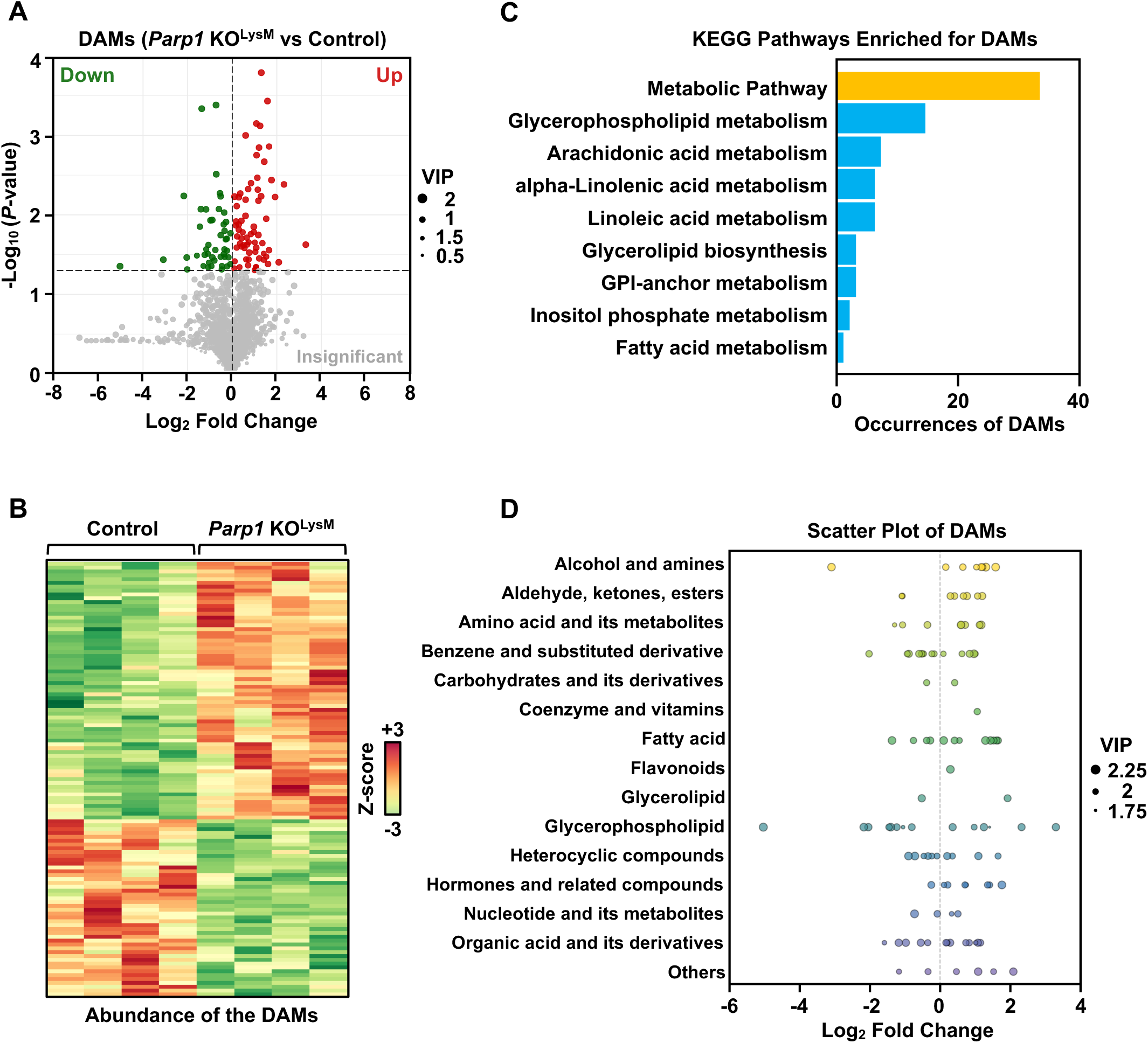
Macrophage PARP1 alters the metabolite profile in WAT. Untargeted metabolomics analysis in WAT from control and *Parp1* KO^LysM^ mice after 4 weeks of HFD feeding (n = 4 for each group). **(A)** Volcano plot showing all the detected metabolites. Each dot in the volcano plot represents a metabolite, where red dots represent upregulated differential metabolites, green dots represent downregulated differential metabolites, and grey dots represent metabolites detected but with insignificant differences. The horizontal coordinate represents changes in the relative metabolite level between two groups [shown as Log_2_ (fold change)], the vertical coordinate indicates the significance level of difference [shown as –Log_10_ (p-value)], and the size of the dot represents the VIP (variable important in projection) score. **(B)** Heatmap of hierarchical clustering showing differentially accumulated metabolites in WAT from *Parp1* KO^LysM^ versus control mice. Each row represents one metabolite, and each column represents one sample. The relative metabolite level is depicted according to the color scale. **(C)** Bar plot showing KEGG pathway analysis of differentially accumulated metabolites in *Parp1* KO^LysM^ versus control mice. **(D)** Category scatter plot showing classification of differentially accumulated metabolites. Each dot represents a metabolite. The horizontal coordinate represents changes in the relative metabolite level between two groups [shown as Log_2_ (fold change)], the vertical coordinate indicates individual category in different color, and the size of the dot represents the VIP (variable important in projection) score.

### Macrophage PARP1 depletion promotes adipose expansion and macrophage accumulation

To determine the effect of macrophage PARP1 on the regulation of adipose tissue functions, we collected both inguinal WAT (iWAT) and epididymal WAT (eWAT) (Figure 4A). The former is recognized as subcutaneous WAT and the latter is recognized as visceral WAT [28]. The weight of both iWAT and eWAT from *Parp1* KO^LysM^ mice was significantly elevated compared to control mice (Figure 4B). H&E staining of eWAT sections, but not iWAT sections, showed a significant increase in adipocyte size in *Parp1* KO^LysM^ mice compared to control mice. Immunofluorescent staining of eWAT for perilipin, a protein located on the surface of intracellular lipid droplets, also demonstrated enlarged adipocytes in *Parp1* KO^LysM^ mice (Supplementary Figure 2A). These results suggests that macrophage-specific depletion of PARP1 promotes WAT expansion and enhances adipocyte hypertrophy in eWAT.

**Figure 4.**
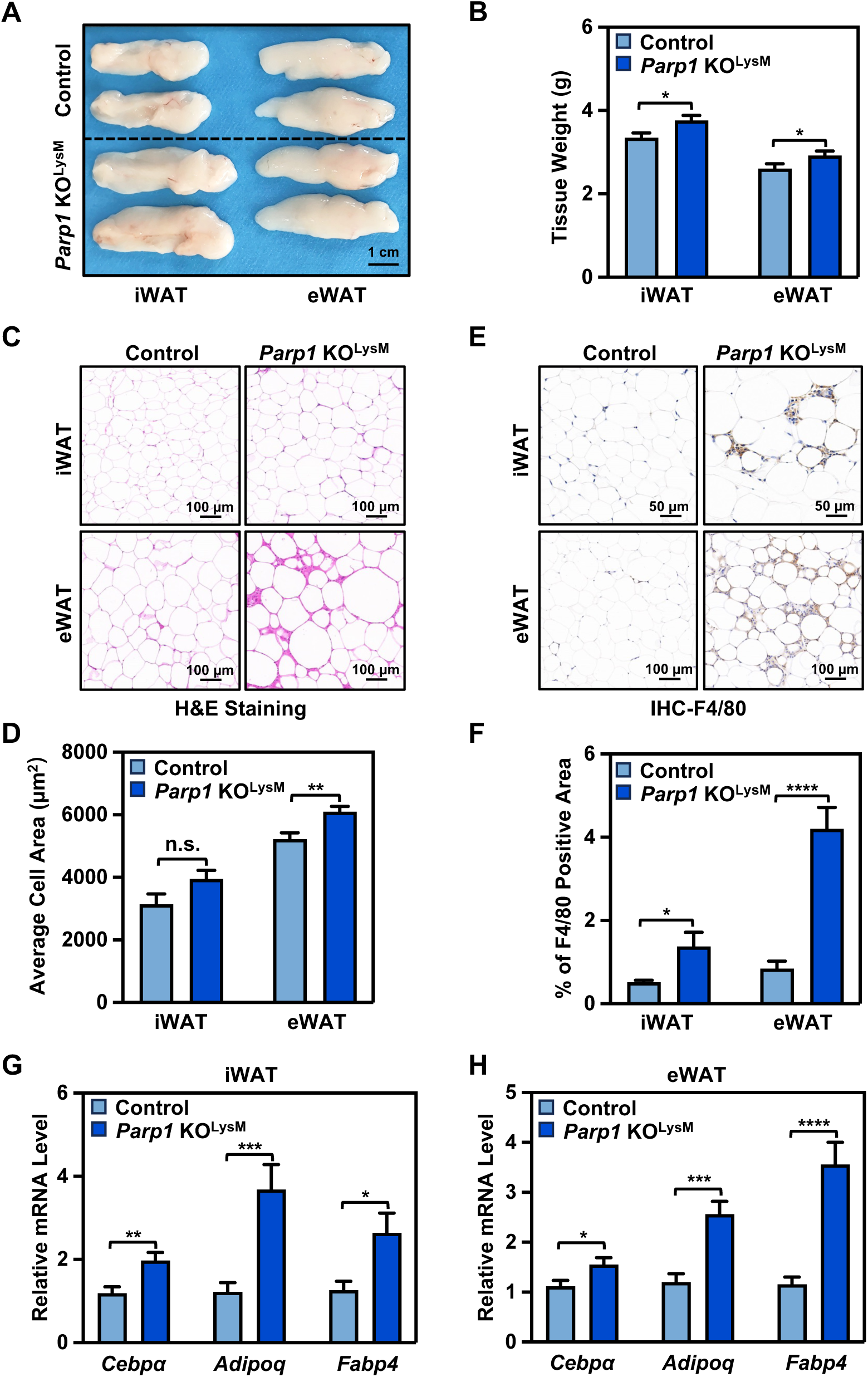
Macrophage PARP1 depletion promotes adipose expansion and macrophage accumulation. **(A)** Representative images of iWAT (*Left*) and eWAT (*Right*) from control and *Parp1* KO^LysM^ mice after 12 weeks of HFD feeding. **(B)** Weight of iWAT and eWAT from control and *Parp1* KO^LysM^ mice after 12 weeks of HFD feeding. Each bar represents the mean + SEM (n = 15 for each group). Asterisks indicate significant differences from the control; two tailed, unpaired t-test; *, p < 0.05. **(C)** H&E staining of iWAT (*Upper panels*) and eWAT (*Lower panels*) from control and *Parp1* KO^LysM^ mice. Scale bar = 100 µm. **(D)** Quantification of H&E staining shown in **(C)**, by analyzing average cell area using ImageJ. Each bar represents the mean + SEM (iWAT, n = 8 for each group; eWAT, n = 11 for each group). Asterisks indicate significant differences from the control; two tailed, unpaired t-test; **, p < 0.01; n.s., no significant difference. **(E)** Immunohistochemical staining of F4/80 for iWAT (*Upper panels*) and eWAT (*Lower panels*) from control and *Parp1* KO^LysM^ mice. Scale bar = 50 µm / 100 µm for iWAT / eWAT as indicated. **(F)** Quantification of immunohistochemical staining shown in **(E)**, by analyzing percent of F4/80 positive area using ImageJ. Each bar represents the mean + SEM (n = 9 for each group). Asterisks indicate significant differences from the control; two tailed, unpaired t-test; *, p < 0.05; and ****, p < 0.0001. **(G and H)** Expression of adipogenic genes *Cebpα*, *Fabp4*, and *Adipq* in iWAT **(G)** and eWAT **(H)** from control and *Parp1* KO^LysM^ mice, as determined by RT-qPCR. Each bar represents the mean + SEM (n = 16 for each group). Asterisks indicate significant differences from the control; two tailed, unpaired t-test; *, p < 0.05; **, p < 0.01; ***, p < 0.001; and ****, p < 0.0001.

Adipose tissue is composed of adipocytes, the stromal vascular fraction (SVF), and extracellular matrix. The SVF derived from the WAT contains heterogeneous cell populations, including preadipocytes (i.e., adipocyte precursors), a population of immune cells, and other diverse cell types, such as stem cells, fibroblasts, endothelial cells, and vascular smooth muscle cells (Figure 1B) [2]. Macrophage accumulation in adipose tissue is an important indication of dysfunctional adipose tissue [29]. To investigate whether macrophage PARP1 affects macrophage infiltration and accumulation in WAT, we performed immunohistochemical staining for F4/80, a well-established macrophage marker, on both iWAT and eWAT sections [30]. The results demonstrate that knockout of *Parp1* in macrophages promoted macrophage infiltration and accumulation in WAT of *Parp1* KO^LysM^ compared to control mice (Figure 4, E and F).

In order to store the excess energy, WAT may expand through two distinct mechanisms: (1) adipocyte hypertrophy, as described above (Figure 4, C and D), and (2) adipocyte hyperplasia (i.e., adipogenesis). We analyzed the expression of adipogenic genes in differentiated adipocytes. We found that *Parp1* KO^LysM^ mice exhibited higher expression of *Cebpα*, *Adipq*, and *Fabp4* in both iWAT (Figure 4G) and eWAT (Figure 4H) compared to control mice, suggesting that enhanced adipogenesis might occur in the WAT of *Parp1* KO^LysM^ mice. In addition, to investigate whether macrophage PARP1 depletion could also affect brown adipose tissue (BAT), we analyzed gene expression in BAT and observed no significant changes in the expression of thermogenic genes, including *Prdm16, Ppargc1a,* and *Ucp1* (Supplementary Figure 2B).

### Depletion of PARP1 in macrophages enhances adipogenesis in WAT

Obesity predisposes a proinflammatory state via increased inflammatory mediators, such as IL-1β, TNF-α, and IL-6 [31, 32]. Adipose tissue macrophages are one of the key immune cells contributing to obesity-associated chronic inflammation [4, 33, 34]. We explored how PARP1 regulates the inflammatory state of macrophages to influence adipocyte differentiation. We isolated SVF from eWAT and sorted macrophages (i.e., F4/80^+^/CD45^+^/CD11b^+^ cells) [30, 35] by flow cytometry. RT-qPCR showed that *Parp1* mRNA expression was significantly decreased in the macrophage population, but not in the total SVF cell population in *Parp1* KO^LysM^ mice compared to control mice (Figure 5A). These results confirmed that the *Parp1* gene was specifically knocked out in macrophages which originate from the myeloid lineage. Depletion of PARP1 in macrophages dramatically downregulated the expression of proinflammatory genes, including *Il1b, Il6,* and *Tnf* (Figure 5B). The expression of these genes was also decreased in the total SVF cell population in *Parp1* KO^LysM^ versus control mice (Supplementary Figure 3A). These results suggest that knockout of *Parp1* in the myeloid lineage specifically impacts the macrophage population in WAT, subsequently alleviating macrophage inflammatory responses to diet-induced obesity.

**Figure 5.**
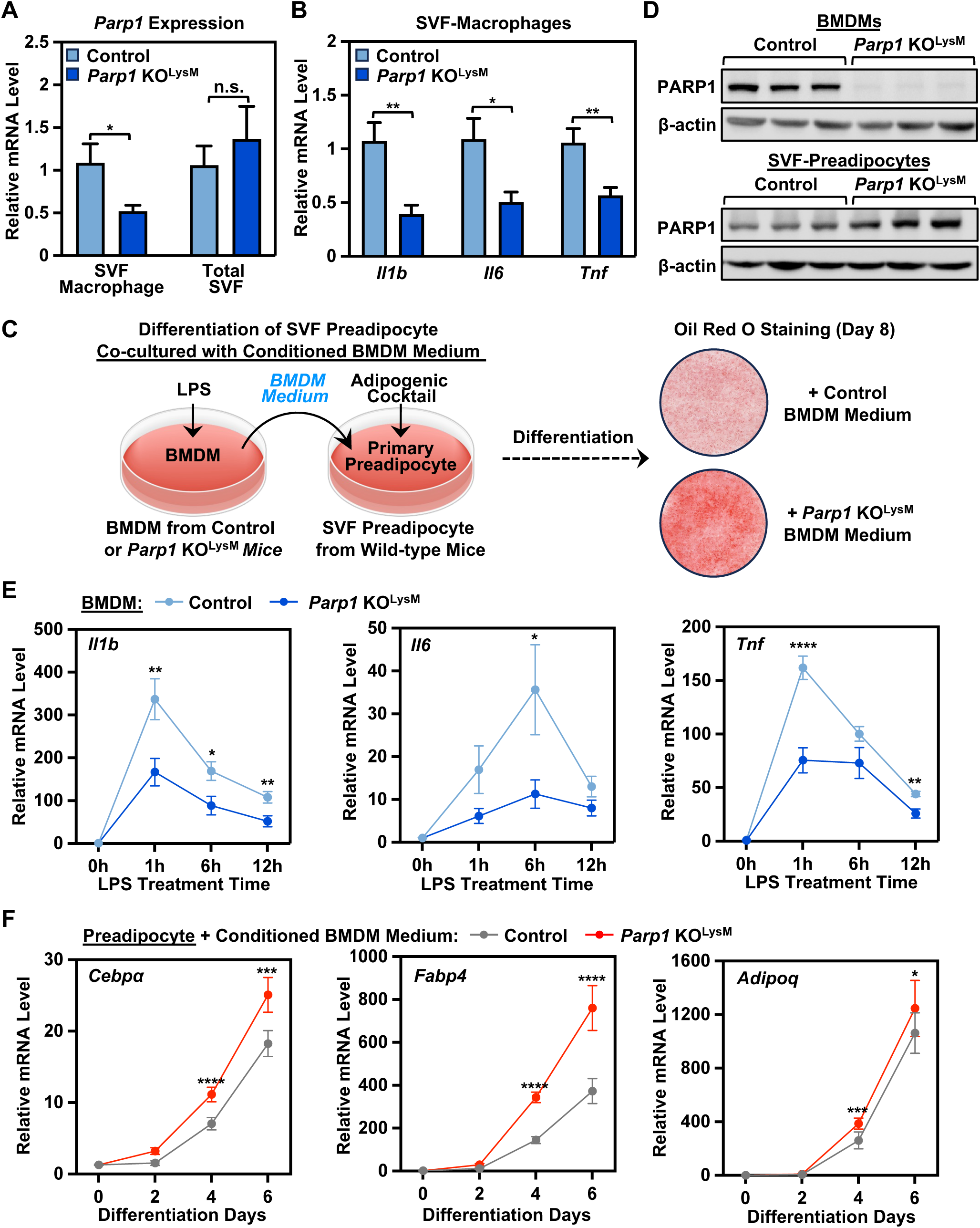
Depletion of PARP1 in macrophages promotes adipogenesis by inhibiting inflammatory responses. **(A)** Expression of *Parp1* gene in sorted macrophage (F4/80^+^/CD45^+^/CD11b^+^) from SVF versus total SVF isolated from eWAT of control and *Parp1* KO^LysM^ mice, as determined by RT-qPCR. Each bar represents the mean + SEM (n = 4 for each group). Asterisks indicate significant differences from the control; two tailed, unpaired t-test; *, p < 0.05; n.s., no significant difference. **(B)** Expression of proinflammatory genes *Il1b, Il6,* and *Tnf* in sorted macrophage (F4/80^+^/CD45^+^/CD11b^+^) from SVF isolated from eWAT of control and *Parp1* KO^LysM^ mice, as determined by RT-qPCR. Each bar represents the mean + SEM (n = 7 for each group). Asterisks indicate significant differences from the control; two tailed, unpaired t-test; *, p < 0.05; and **, p < 0.01. **(C)** Schematic representation of differentiation of SVF preadipocytes from wild-type C57BL/6 mice using the conditioned medium from LPS-stimulated BMDMs isolated from control or *Parp1* KO^LysM^ mice. Oil red O staining showing representative images of lipid droplet formation in the differentiated adipocytes at differentiation day 8. **(D)** Western blotting showing PARP1 expression in BMDMs and SVF-preadipocytes from control or *Parp1* KO^LysM^ mice (n = 3 for each group). β-actin was used as loading control. **(E)** Expression of proinflammatory genes *Il1b, Il6,* and *Tnf* in BMDMs isolated from control or *Parp1* KO^LysM^ mice, upon the LPS treatment, as determined by RT-qPCR. Each point represents the mean ± SEM (*Il1b*, n = 16 for each group*; Il6,* n = 10 for each group*; Tnf,* n = 12 for each group). Asterisks indicate significant differences from the control at individual time points; two tailed, unpaired t-test; *, p < 0.05; **, p < 0.01; ***, p < 0.001; and ****, p < 0.0001. **(F)** Expression of adipogenic genes *Cebpα*, *Fabp4*, and *Adipq* in wild-type SVF preadipocytes induced differentiating with culture medium containing adipogenic cocktail and conditioned medium from LPS-stimulated BMDMs isolated from control or *Parp1* KO^LysM^ mice, as determined by RT-qPCR. Each bar represents the mean ± SEM (*Cebpα* and *Fabp4,* n = 9 for each group*; Adipq,* n = 11 for each group). Asterisks indicate significant differences from the control at individual time points; two tailed, unpaired t-test; *, p < 0.05; ***, p < 0.001; and ****, p < 0.0001.

To investigate whether macrophage PARP1 depletion could directly affect adipogenesis through decreased proinflammatory cytokine secretion, we used the conditioned medium from bone marrow-derived macrophages (BMDMs) isolated from *Parp1* KO^LysM^ or control mice, to culture and differentiate SVF preadipocytes isolated from wild-type C57BL/6 mice (Figure 5C). To generate the conditioned medium, BMDMs from *Parp1* KO^LysM^ or control mice were treated with lipopolysaccharide (LPS) to stimulate an inflammatory response reminiscent of the obese adipose tissue microenvironment [36, 37]. We first examined PARP1 expression in cultured BMDMs and SVF preadipocytes. Western blotting demonstrated that PARP1 expression was lost in BMDMs, while remained unchanged in SVF preadipocytes isolated from *Parp1* KO^LysM^ mice, compared to control mice (Figure 5D). RT-qPCR also showed that *Parp1* mRNA expression was barely detectable in BMDMs from *Parp1* KO^LysM^ mice (Supplementary Figure 3B), confirming the successful knockout of the *Parp1* gene in macrophages in our transgenic mouse model.

Treatment of BMDMs from control mice with LPS induced the expression of proinflammatory cytokines, including *Il1b, Il6,* and *Tnf* (Figure 5E). The expression of these genes reached the peak between 1 hour (*Il1b* and *Tnf*) and 6 hours (*Il6*) after LPS treatment and declined after 12 hours of treatment (Figure 5E). Compared to BMDMs from control mice, BMDMs from *Parp1* KO^LysM^ mice demonstrated significantly decreased expression of *Il1b, Il6,* and *Tnf* (Figure 5E), indicating that depletion of PARP1 inhibits the LPS-induced proinflammatory response in macrophages. We then induced differentiation of primary SVF preadipocytes isolated from wild-type C57BL/6 mice and cocultured them with the conditioned medium of LPS-treated BMDMs from control or *Parp1* KO^LysM^ mice respectively, throughout 8 days of differentiation. Exposure of the preadipocytes to the conditioned BMDM medium from *Parp1* KO^LysM^ mice: (1) enhanced lipid droplet formation in mature adipocytes compared to control mice, as detected by oil red O staining (Figure 5C) and (2) significantly increased the expression of proadipogenic genes, including *Cebpα*, *Fabp4*, and *Adipq* (Figure 5F). Taken together, these results demonstrate that depletion of PARP1 in macrophages promotes adipogenesis by downregulating inflammatory responses.

### Depletion of PARP1 in macrophages alters macrophage gene expression and the cellular composition of adipose tissue

To further dissect the inflammatory signature regulated by PARP1 depletion in the myeloid lineage, we performed single cell RNA-sequencing (scRNA-seq) using sorted CD45^+^ (a pan-leukocyte marker) immune cells from SVF of WAT collected from *Parp1* KO^LysM^ versus control mice [38]. Clustering of the data from 64,257 cells obtained from three replicates of control mice (35,385 cells) and *Parp1* KO^LysM^ mice (28,872 cells) revealed 17 distinct cell populations (Figure 6A), indicating the heterogeneity of immune cells in WAT. For better cell type identification, we used established marker genes to classify these cells into 8 populations, including T cells, B cells, monocytes, macrophages, NK cells, neutrophils, endothelial-like immune cells, and fibroblast-like immune cells (Figure 6B), each containing cells from both control and *Parp1* KO^LysM^ mice (Figure 6C).

**Figure 6.**
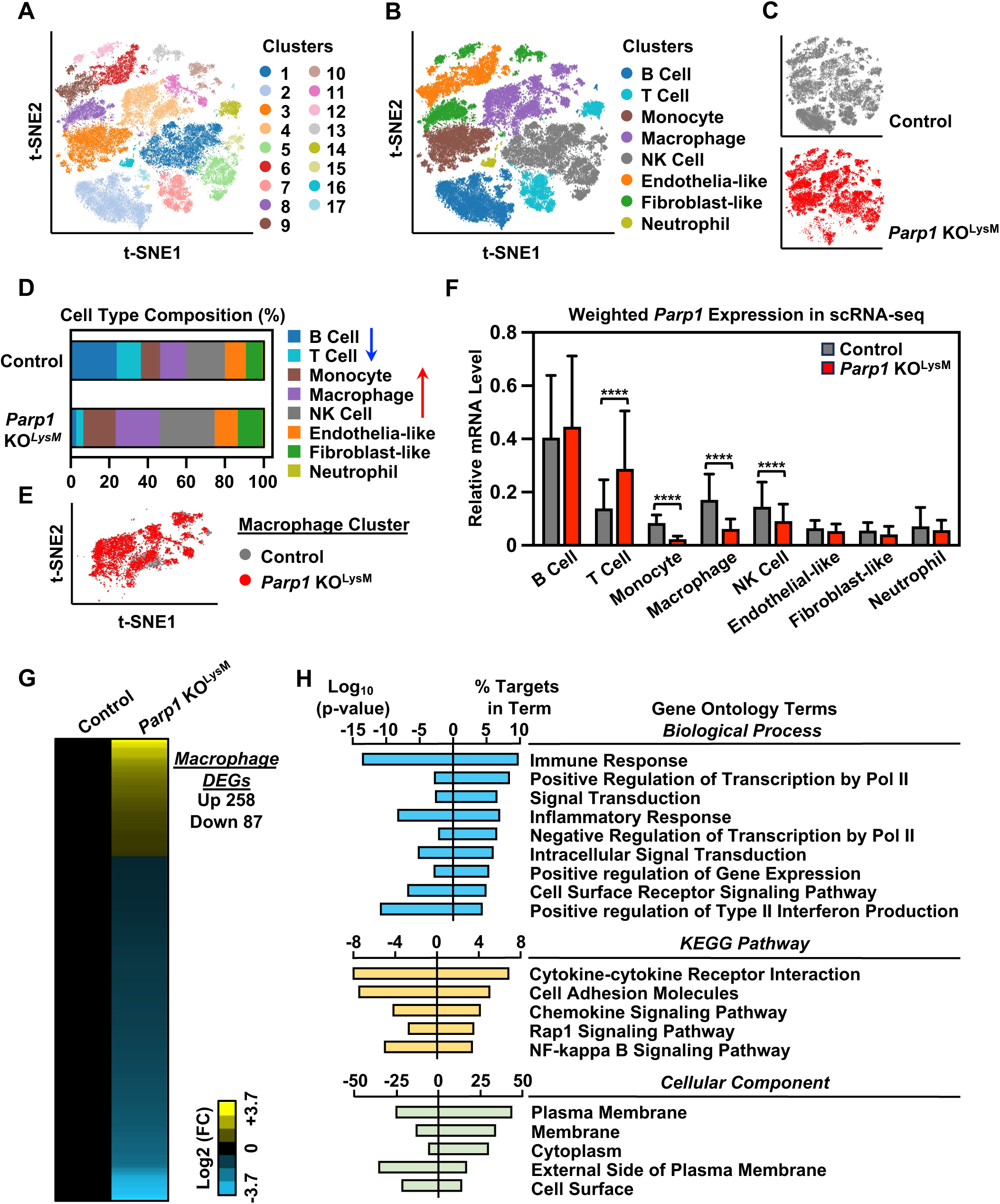
Depletion of PARP1 in macrophages alters the cellular composition and macrophage gene expression in WAT. scRNA-seq analysis of sorted CD45^+^ immune cells from SVF of WAT in control and *Parp1* KO^LysM^ mice after 12 weeks of HFD feeding (n = 3 for each group). **(A)** Identification of 17 different cell clusters from 64257 cells obtained from control mice (35385) and *Parp1* KO^LysM^ mice (28872) analyzed by t-SNE clustering approach. Each point depicts a cell, colored according to cluster designation. **(B)** t-SNE plot showing annotation of cell clusters using established marker genes. Each point depicts a cell, colored according to cluster designation. **(C)** t-SNE plot showing cell clusters annotated to control (grey) and *Parp1* KO^LysM^ (red) groups. Each point depicts a cell. **(D)** Bar plots showing percent of 8 identified cell subpopulations in control and *Parp1* KO^LysM^ groups, respectively, colored according to cluster designation. **(E)** t-SNE plot showing macrophage cluster annotated to control (grey) and *Parp1* KO^LysM^ (red) groups. Each point depicts a cell. **(F)** Bar plots of expression level of *Parp1* gene in 8 identified cell subpopulations between control and *Parp1* KO^LysM^ groups from scRNA-seq datasets. Each bar represents the mean + SEM. Asterisks indicate significant differences from the control in individual cell clusters; Wilcoxon Signed-Rank test; ****, p < 2.2 × 10^-16^. **(G)** Heatmaps showing differentially expressed genes (DEGs) in macrophage cluster between control and *Parp1* KO^LysM^ groups from scRNA-seq. **(H)** Gene Ontology terms for DEGs in macrophage cluster shown in **(G)**.

We observed an increase in the fraction of macrophages among the immune cell populations in *Parp1* KO^LysM^ mice (∼22.2%) compared to control mice (∼13.1%) (Figure 6, D and E). The monocyte and NK cell fractions also increased in *Parp1* KO^LysM^ mice (Figure 6D). In contrast, the fractions of T cells and B cells were decreased in *Parp1* KO^LysM^ mice (Figure 6D). Since the *Parp1* gene was specifically deleted in the myeloid lineage in our mouse model, we compared *Parp1* gene expression in different cell clusters between control and *Parp1* KO^LysM^ mice. As expected, *Parp1* gene expression was significantly decreased in macrophages and monocytes, both of which originate from myeloid lineage, in *Parp1* KO^LysM^ mice compared to control mice (Figure 6F). Interestingly, we also observed a dramatic reduction in *Parp1* expression in NK cells, which mainly originate from the lymphoid lineage (Figure 6F). In contrast, *Parp1* expression in T cells was upregulated in *Parp1* KO^LysM^ mice versus control mice and remained unchanged in other cell clusters (Figure 6F). These results suggest that deletion of PARP1 in the myeloid lineage leads to a major alteration of the cellular composition in WAT.

To further investigate the effects of macrophage PARP1 on gene expression, we analyzed the differentially expressed genes (DEGs) in macrophages from *Parp1* KO^LysM^ mice versus control mice. We observed a total of 345 genes that exhibited significantly altered expression (Figure 6G), which were enriched in immune/inflammatory responses, transcription regulation, signal transduction, and cell surface receptor signaling pathway (Figure 6H). KEGG pathway analysis demonstrated an enrichment in cytokine-cytokine receptor interactions, and cellular component analysis showed cell surface and membrane localization of most of these genes (Figure 6H). These analyses suggest that the regulatory effects of macrophage PARP1 on WAT are mediated through interactions between macrophages and other immune cell populations.

### Depletion of PARP1 in macrophages enhances crosstalk between macrophages and NK cells

To identify potential ligand–receptor interactions between immune cell populations in an unbiased manner, we performed cell communication analysis using CellChat v1.6 [39]. Chord diagrams demonstrated that the signaling interactions between NK cell and macrophage were significantly enhanced, while the interactions between T cell and B cell were lost, in *Parp1* KO^LysM^ mice compared to control mice (Figure 7A). Bubble plots revealed that the *Ptprc* (gene encoding a ligand on NK cells)–*Mrc1* (gene encoding an activating receptor on macrophages) interactions were significantly upregulated in WAT immune cell populations in *Parp1* KO^LysM^ mice versus control mice (Figure 7B).

**Figure 7.**
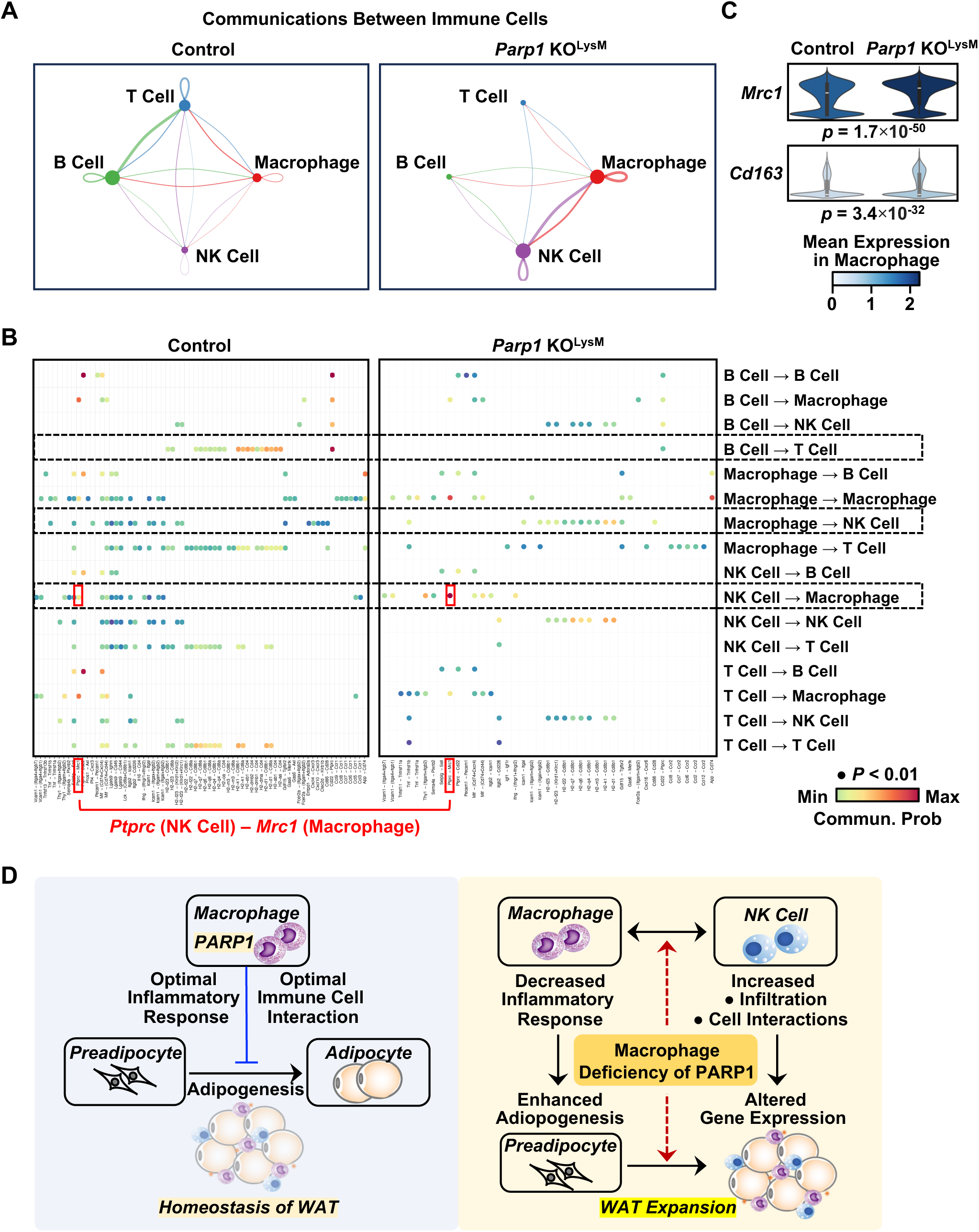
Depletion of PARP1 in macrophages enhances communication between macrophages and NK cells. **(A)** Chord diagrams showing the signaling interactions among cell subpopulations from scRNA-seq (macrophage, NK cell, T cell, and B cell) between control and *Parp1* KO^LysM^ groups, analyzed by CellChat. **(B)** Bubble plots showing ligand-receptor interaction pathways among cell subpopulations from scRNA-seq between control and *Parp1* KO^LysM^ groups, analyzed by CellChat. **(C)** Violin plots showing expression level of M2-type marker genes *Mrc1* and *Cd136* in macrophage cluster from scRNA-seq between control and *Parp1* KO^LysM^ groups, analyzed by Wilcoxon Signed-Rank test (p value as indicated). **(D)** Schematic presentation showing that macrophage PARP1 is critical in the maintenance of adipose tissue inflammatory and metabolic homeostasis. Deficiency of PARP1 in macrophage not only affect macrophage, but also promotes cell-cell crosstalk among macrophage, preadipocyte, adipocyte and NK cell, driving the development of obesity.

MRC1, also known as CD206, is a well-established marker for alternative activation of macrophages (M2-type) in both humans and mice [40, 41]. We found that the expression of *Mrc1* as well as *Cd163*, another M2-type macrophage marker, was significantly upregulated in macrophage cluster from *Parp1* KO^LysM^ mice compared to control mice (Figure 7C). In contrast, *Cd68* and *Cd80*, two M1-type macrophage markers remained unchanged or mildly changed in macrophages between control and *Parp1* KO^LysM^ mice (Supplementary Figure 4A). In addition, the fraction of cells expressing *Mrc1* or *Cd163* within the macrophage cluster was higher in *Parp1* KO^LysM^ mice than in control mice (Supplementary Figure 4B). These results suggests that macrophage PARP1 depletion drives a phenotypic switch of macrophages towards an M2-like state, consistent with the downregulation of proinflammatory gene expression in macrophages from WAT of *Parp1* KO^LysM^ mice shown in Figure 5B.

In addition to adipose tissue macrophages, alterations in NK cells also contribute to obesity and obesity-related metabolic outcomes [42]. We showed that macrophage PARP1 depletion increased the NK cell population in adipose tissue (Figure 6D) and enhanced cell communications between NK cells and macrophages (Figure 7A), as described above. Here, we further examined the regulatory effects of macrophage PARP1 on gene expression in NK cells from our scRNA-seq data. We observed a total of 1,587 genes differentially expressed in NK cells from the WAT of *Parp1* KO^LysM^ mice compared to control mice, the majority of which (1,477 genes) were downregulated (Supplementary Figure 4C). Interestingly, KEGG analysis demonstrated that these genes were primarily enriched in metabolic pathways (Supplementary Figure 4D), suggesting an important role of NK cells in regulating metabolism in WAT. Taken together, our results demonstrate that macrophage-specific PARP1 depletion alters the cell composition and transcriptional profiles of WAT during diet-induced obesity, in particular by enhancing cell-cell crosstalk between macrophages and NK cells.

## Discussion

In the studies described herein, we documented a previously unrecognized role for macrophage PARP1 in regulating the differentiation of preadipocytes and promoting interactions between immune cells within adipose tissue. These results provide evidence that macrophage PARP1 is required to maintain an optimal inflammatory state and metabolic homeostasis in white adipose tissue. Depletion of PARP1 in macrophages, on the other hand, (1) disrupts inflammatory responses in macrophages, (2) enhances adipogenesis and adipocyte expansion, and (3) promotes infiltration and interactions of macrophages and NK cells, thus exacerbating the development of diet-induced obesity and related metabolic pathologies, such as fatty liver disease (Figure 7D).

### Biological functions of PARP1 in metabolic physiology and disorders

Historically, PARP1 has been linked to DNA damage responses and cancers. Nonetheless, a growing body of evidence has shed new light on biological functions of PARP1 in a broad array of physiological and pathological processes [7]. Numerous studies over the past decades have connected PARP1 catalytic activity to metabolic outcomes, including obesity, hyperlipidemia, type II diabetes, and fatty liver diseases [11–21, 43–47]. Genetic deletion of PARP1 or pharmacological inhibition of PARP1 activity can have major impacts on metabolism, including obesity, lipid metabolism, glucose-insulin metabolism, and fatty liver disease [11–21, 43–47]. However, the metabolic phenotypes of whole-body *Parp1* knockout mice have not always been consistent [13, 44, 48–50]. This is likely due to the fact that PARP1 is widely expressed across metabolic organs and tissues, including adipose tissue, liver, skeletal muscle, pancreas, brain and the nervous system [22], and effects in one tissue may lead to effects in another.

In this regard, developing and using tissue-specific *Parp1* knockout genetic mouse model has helped achieve a better understanding of functional roles of PARP1 in metabolic physiology and disorders. To unambiguously address these questions, we have recently examined the role of tissue-specific PARP1 in the control of adipogenesis using a preadipocyte-specific conditional *Parp1* knockout with lineage-tracing mouse genetic model [11]. Our results demonstrated a definitive role for PARP1 in repressing the expansion of adipocyte precursor populations to limit adipogenesis, consistent with previous observations [17, 20, 44]. In this study, we further explored the role of macrophage PARP1 in adipogenesis and obesity, by using a myeloid lineage-specific *Parp1* knockout mouse genetic model. Our results indicates that depletion of PARP1 in macrophages enhances the differentiation of preadipocyte via distinct mechanisms. Thus, PARP1 acts simultaneously in both adipose tissue macrophages and adipocyte precursors to control the adipocyte population and adipose tissue expansion, promoting a “healthy” fat environment to limit adverse metabolic consequences. Similar to the metabolic phenotype of preadipocyte-*Parp1* knockout, macrophage-*Parp1* knockout promoted diet-induced obesity, and further the development of NAFLD in mice. We previously reported that pharmacological inhibition of PARP1 activity or whole-body knockout of *Parp1* alleviated diet-induced hepatic steatosis and inflammation [16]. In this regard, the observation of NAFLD development in macrophage-specific *Parp1* knockout mice reemphasizes the fact that metabolic effects of PARP1 are mediated in a tissue-specific manner. Both adipose tissue macrophage and hepatic macrophage may contribute to the obesity-associated NAFLD in this tissue-specific *Parp1* knockout mouse model.

### Adipose tissue macrophages modulate the inflammatory state in obesity: Role of PARP1

Adipose tissue contains a diverse population of immune cells, among which macrophages play a critical role in maintaining adipose tissue homeostasis and promoting low-grade inflammation during the development of obesity [33, 51]. Adipose tissue expansion is tightly associated with (1) the accumulation of macrophages in adipose tissue and (2) the phenotypic switch of macrophage polarization [33, 51, 52]. The increased number of macrophages in obese adipose tissue could originate from the recruitment of monocyte-derived macrophages [51], or local proliferation of tissue-resident macrophages [53]. However, it is difficult to distinguish them by cell surface markers. In this study, we demonstrated that depletion of PARP1 in macrophages enhanced macrophage accumulation, and promoted a phenotypic switch of macrophage polarization in the obese adipose tissues.

Macrophages can be classified into two major populations: classically activated M1-type macrophages and alternatively activated M2-type macrophages. However, emerging evidence has identified a broader range of macrophage populations than the classical M1/M2 model [54]. Although preclinical studies showed that obesity predisposes a proinflammatory state of macrophages [33, 52], some human studies demonstrated the phenotypic shift of adipose tissue macrophages towards a more anti-inflammatory cell type in obese subjects [55, 56], suggesting that the common paradigm may not reflect the whole picture in human obesity. Our results suggest that macrophage depletion of PARP1 leads to a phenotypic switch towards a more M2-type like macrophage, with increased expression of M2 cell surface markers (*Mrc1* and *Cd163*), as well as suppressed inflammatory responses. The expression of M1-type macrophage markers was not correspondingly decreased upon macrophage depletion of PARP1, indicating the complex range of macrophage subtypes and activation states in the adipose tissue. In addition, the metabolite profile may also affect macrophage phenotype [57, 58]. The differentially accumulated metabolites in WAT of *Parp1* KO^LysM^ in our metabolomics analysis are mostly involved in glycerophospholipid and polyunsaturated fatty acid metabolism, which may contribute to polarization of macrophages to an anti-inflammatory state. Taken together, our studies provide evidence that macrophage PARP1 is required to maintain an “optimal” inflammatory state in macrophages and adipose tissue to create a microenvironment for tissue homeostasis.

### Regulation of cell-cell crosstalk within white adipose tissue by macrophage PARP1

Besides adipocytes, adipose tissue contains a heterogeneous cell population, including adipocyte precursors, immune cells, and other cell types [2]. All these cell types can contribute both separately and cooperatively to maintain adipose tissue homeostasis. However, it is unclear how different cell types cooperate through cell-cell crosstalk within adipose tissue. To address this question, we investigated the role of macrophage-dependent PARP1 in regulating cell communications among (1) macrophage, preadipocyte and adipocyte; and (2) immune cell populations. Obesity is often characterized by increases in both the size of adipocyte (hypertrophy) and the number of newly synthesized adipocyte (adipogenesis). Hypertrophic adipocytes can secrete a variety of adipokines, which contribute importantly to obesity-related complications, such as type II diabetes, and NAFLD [59, 60]. Our studies showed that depletion of PARP1 in macrophages not only enhances adipocyte hypertrophy, but also promotes adipogenesis through downregulation of inflammatory responses. These results are consistent with previous studies, which have demonstrated that proinflammatory macrophages impairs adipogenesis [61, 62]. Therefore, macrophage PARP1 optimizes the inflammatory state of adipose tissue to control the adipocyte population in a metabolic healthy state. Depletion of PARP1 in macrophages decreases macrophage inflammatory responses, and then drives metabolic dysfunction of adipose tissue through the crosstalk between macrophages and adipocytes.

Adipose tissue contains a wide range of immune cell populations. Depletion of PARP1 in macrophages affects immune cell composition in adipose tissue, involving macrophages, monocytes, NK cells, T cells, and B cells. Emerging evidence demonstrated that NK cell, another key regulator in adipose tissue to drive obesity and insulin resistance, has a significant impact on macrophages [42, 63, 64]. Depletion of PARP1 in macrophages increased infiltration of both macrophages and NK cells in adipose tissue, altered gene expression, and enhanced the interactions between these two cell populations. Thus, our studies revealed a critical role of macrophage-dependent PARP1 in regulating crosstalk among immune cell populations. In addition, previous studies have shown that PARP1 expression and activity can regulate the activation of NK cells and T cells [65–67]. In our study, the changes of PARP1 expression in NK cells and T cells upon macrophage PARP1 depletion might contribute to the dysfunction of both cell types in WAT during obesity. However, the definitive molecular mechanism underlying immune cell interactions needs further investigation.

Collectively, our study has identified a key role of macrophage PARP1 in the maintenance of adipose tissue inflammatory and metabolic homeostasis. Depletion of PARP1 in macrophages not only affects macrophages, but also promotes cell-cell crosstalk among preadipocyte, adipocyte and immune cells, which cooperatively drive the development of obesity and related metabolic outcomes.

## Materials and Methods

### Antibodies and specialized reagents

The following antibodies were used for Western blotting: PARP1 (Cell Signaling, 9532; RRID:AB_659884), β-actin (Abcam, ab8226; RRID:AB_306371), goat anti-rabbit HRP-conjugated IgG (ThermoFisher, 31460; RRID:AB_228341), and goat anti-mouse HRP-conjugated IgG (ThermoFisher, 31430; RRID:AB_228307). The following antibodies were used for immunohistochemistry and immunofluorescent: F4/80 (Cell Signaling, 70076; RRID: AB_2799771), perilipin (Fitzgerald, 20R-PP004; RRID:AB_1288416), goat anti-rabbit biotin-conjugated IgG (ThermoFisher, 65-6140, RRID:AB_2533969), and Alexa fluor 647-conjugated goat anti-guinea pig IgG (ThermoFisher, A-21450; RRID:AB_2535867). The following antibodies were used for flow cytometry: CD45-APC/Cy7 (BD Biosciences, 557659; RRID:AB_396774), CD11b-FITC (BD Bioscience, 557396; RRID:AB_396679), and F4/80-BV421 (BD Bioscience, 565411; RRID:AB_2734779).

Other chemicals for cell treatments included: LPS (Sigma-Aldrich, L2630), M-CSF (GenScript, Z02930), insulin (Biosharp, BS901), 3-isobutyl-1-methylxanthine (Sigma, 410957), dexamethasone (Sigma, D4902), and indomethacin (Sigma-Aldrich, I7378).

### Mouse studies

All animal experiments were performed in compliance with the Institutional Animal Care and Use Committee (IACUC) of UT Southwestern Medical Center, and IACUC of Tongji Medical College of Huazhong University of Science and Technology, respectively. All animal procedures were conducted according to NIH Guide for the Care and Use of Laboratory Animals. *Parp1^loxP/loxP^*mice were generated in the Kraus Laboratory from the European Conditional Mouse Mutagenesis Program stock [20] and are available from the Jackson Laboratory (stock No. 032650). *LysM-Cre* mice were described previously [68], and purchased from the Jackson Laboratory (stock No. 004781). *Parp1^loxp/loxp^* mice were crossed with *LysM-Cre* mice to generate myeloid lineage-specific *Parp1* knockout mice (*Parp1* KO^LysM^, i.e., *Parp1 ^loxp/loxp^*/*LysM-Cre^tg/-^*) (Figure 1A), and their *Parp1^loxp/loxp^*/*LysM-Cre^-/-^* littermates were used as control. Mice were maintained in the animal facility on a 12-hour light and dark cycle with free access to food and water.

***High-fat diet (HFD) treatment.*** Eight-week-old male *Parp1* KO^LysM^ and control mice were switched from standard chow diet to HFD (60 kcal% fat; Research Diets, D12492) for 12 weeks. Body weight of the mice was monitored weekly. Body composition of the mice was examined using an EchoMRI-100 Body Composition Analyzer at UT Southwestern Metabolic Phenotyping Core every two weeks. At the end of the experiment, fresh tissues were collected from the mice, rapidly frozen in liquid nitrogen, and stored at –80°C until used.

### Metabolic chamber analyses

Energy metabolism of the HFD-fed mice described above was monitored using the Columbus Comprehensive Lab Animal Monitoring System (CLAMS, Columbus. Instruments, Columbus, OH) through the measurement of oxygen (O_2_) consumption, carbon dioxide (CO_2_) production, and heat generation.

### Glucose tolerance test (GTT) and insulin tolerance test (ITT)

*Parp1* KO^LysM^ and control mice described above were subjected to GTT and ITT after 12 weeks of HFD feeding. Briefly, GTT was performed by intraperitoneal injection of glucose (2 g/kg body weight; Sigma-Aldrich, D9434) after overnight fasting. Tail blood glucose levels was measured with a glucometer before (0 min) and at 15, 30, 45, 60 and 120 min after glucose administration. ITT was performed by intraperitoneal injection of insulin (0.75 U/kg body weight; Biosharp, BS901) after 4-hour fasting. Tail blood glucose levels was measured with a glucometer before (0 min) and at 15, 30, 45, 60 and 120 min after insulin administration.

### Serum lipid analyses

*Parp1* KO^LysM^ and control mice were fed with HFD for 12 weeks as described above. Serum lipid levels of the mice were measured by using the Wako LabAssay kits (290-63701 for triglyceride, and 294-65801 for cholesterol) according to the manufacturer’s instructions.

### RNA isolation and RT-qPCR

Total RNA from the tissue sample or cultured cells were isolated using Trizol Reagent (Takara Bio, 9109) and reverse transcribed using the PrimeScrip RT Master Mix kit (Takara Bio, RR036A) to generate cDNA according to the manufacturer’s protocol. The cDNA samples were subjected to real-time qPCR with SYBR Green Supermix (Bio-Rad, 1708880) on a Bio-Rad CFX-96 real-time system, using gene-specific primers as listed below. Melting curve analyses were performed to ensure that only the targeted amplicon was amplified. Target gene expression was normalized to the expression of *Rn18s.* Relative gene expression was calculated using the comparative counting method Formula 2^−ΔΔCt^. All experiments were performed at least three times with independent biological replicates to ensure reproducibility. All experimental groups that were compared had similar variance as determined by the standard deviation of the biological replicates within each group. Statistical differences between two groups were determined using the unpaired t-test and a statistical significance was defined as p < 0.05.

***Primers used for RT-qPCR***. The following primers were synthesized by Wuhan Qingke Biotechnology Co., Ltd. (Wuhan, China) and used for RT-qPCR.

*Parp1* forward: 5’-GCTTTATCGAGTGGAGTACGC –3’

*Parp1* reverse: 5’-GGAGGGAGTCCTTGGGAATAC –3’

*Il1* forward: 5’-GAAATGCCACCTTTTGACAGTG –3’

*Il1* reverse: 5’-TGGATGCTCTCATCAGGACAG –3’

*Il6* forward: 5’-CTGCAAGAGACTTCCATCCAG –3’

*Il6* reverse: 5’-AGTGGTATAGACAGGTCTGTTGG –3’

*Tnf* forward: 5’-CAGGCGGTGCCTATGTCTCTC –3’

*Tnf* reverse: 5’-CGATCACCCCGAAGTTCAGTAG –3’

*Fabp4* forward: 5’-AAGGTGAAGAGCATCATAACCCT –3’

*Fabp4* reverse: 5’-TCACGCCTTTCATAACACATTCC –3’

*Adipoq* forward: 5’-TGTTCCTCTTAATCCTGCCCA –3’

*Adipoq* reverse: 5’-CCAACCTGCACAAGTTCCCTT –3’

*Cebpa* forward: 5’-GCGGGAACGCAACAACATC –3’

*Cebpa* reverse: 5’-GTCACTGGTCAACTCCAGCAC –3’

*Srebp1* forward: 5’-TGACCCGGCTATTCCGTGA –3’

*Srebp1* reverse: 5’-CTGGGCTGAGCAATACAGTTC –3’

*Scd1* forward: 5’-TTCTTGCGATACACTCTGGTGC –3’

*Scd1* reverse: 5’-CGGGATTGAATGTTCTTGTCGT –3’

*Fasn* forward: 5’-GGCTCTATGGATTACCCAAGC –3’

*Fasn* reverse: 5’-CCAGTGTTCGTTCCTCGGA –3’

*Prdm16* forward: 5’-CCACCAGCGAGGACTTCAC –3’

*Prdm16* reverse: 5’-GGAGGACTCTCGTAGCTCGAA –3’

*Ppargc1a* forward: 5’-TATGGAGTGACATAGAGTGTGCT –3’

*Ppargc1a* reverse: 5’-GTCGCTACACCACTTCAATCC –3’

*Ucp1* forward: 5’-GTGAACCCGACAACTTCCGAA –3’

*Ucp1* reverse: 5’-TGCCAGGCAAGCTGAAACTC –3’

*Rn18s* forward: 5’-TTGACGGAAGGGCACCACCAG –3’

*Rn18s* reverse: 5’-GCACCACCACCCACGGAATCG –3’

### Untargeted metabolomics

*Parp1* KO^LysM^ and control mice were fed with HFD for 4 weeks. Fresh epididymal white adipose tissue samples were collected from the mice, rapidly frozen in liquid nitrogen, and stored at –80°C until used. For unbiased detection of metabolites, the samples were prepared for untargeted metabolomics. Briefly, the samples were thawed on ice, mixed with extraction solution (ACN:Methanol = 1:4, V/V) containing internal standards, and then centrifuged at 12000 rpm for 10 min at 4 °C. The supernatant was collected and placed in –20 °C for 30 min, and then centrifuged at 12000 rpm for 3 min at 4 °C. The supernatant were collected and analyzed by the LC-MS untargeted metabolomics performed by Metware Biotechnology, Inc., Wuhan, China.

***Data analysis.*** The original data file from LC-MS was converted into mzML format by ProteoWizard software. Peak extraction, peak alignment and retention time correction were performed by XCMS program, respectively. The “SVR” method was used to correct the peak area. The peaks with detection rate lower than 50 % in each group of samples were discarded. Metabolic identification information was obtained by searching the laboratory’s self-built database, integrated public database, AI database and metDNA. Unsupervised Principal component analysis (PCA) was performed by statistics function prcomp within R (www.r-project.org). The data was unit variance scaled before unsupervised PCA. Hierarchical cluster analysis (HCA) was presented as heatmaps with dendrograms, while the Pearson correlation coefficient (PCC) between samples were calculated by the cor function in R and presented as heatmaps. Both HCA and PCC were carried out by R package Complex Heatmap. For HCA, normalized signal intensities of metabolites (unit variance scaling) were visualized as a color spectrum. Differential metabolites were determined by VIP (VIP > 1) and P-value (P-value < 0.05, Student’s t test). VIP values were extracted from OPLS-DA result generated by using R package MetaboAnalystR. In order to avoid overfitting, a permutation test (200 permutations) was performed. KEGG annotation and enrichment analysis was performed to the identified metabolites using KEGG Compound database (http://www.kegg.jp/kegg/compound/), and KEGG Pathway database (http://www.kegg.jp/kegg/ pathway.html). Significantly enriched pathways were identified with a hypergeometric test’s P-value for a given list of metabolites.

### Histological analysis of adipose tissue

Subcutaneous and visceral adipose tissue were collected from *Parp1* KO^LysM^ and control mice described above. Tissue samples were fixed overnight in buffered formaldehyde (10%), embedded in paraffin wax, and then sectioned (5 μM). Sections were mounted on slides, deparaffinized in xylene, and rehydrated in a graded alcohol series. Sections were then stained with hematoxylin and eosin (H&E, Sigma-Aldrich).

***Immunohistochemistry (IHC) and immunofluorescence (IF).*** Briefly, sections were subjected to antigen retrieval by incubating sections in boiling 10 mM sodium citrate buffer (pH 6.0) for 10 min, followed by cooling to room temperature. For peroxidase-based staining, sections were incubated with 3% hydrogen peroxide for 30 min, and then blocked and incubated with primary antibody overnight at 4 °C in 1% bovine serum albumin (BSA) in PBS. After washing with PBS, the slides for IHC were incubated with biotin-conjugated secondary antibody for 1 hour at room temperature, developed by DAB (3,3′ Diaminobenzidine Tetrahydrochloride) solution (Sigma-Aldrich, D7679), and counter-stained with hematoxylin for nuclei. The slides for IF were incubated with Alexa fluor-conjugated secondary antibody for 1 hour at room temperature. The coverslips were then mounted with Anti-Fade mounting medium with DAPI (Vector Laboratories, H-1200). IF images were acquired using an inverted Zeiss LSM 780 confocal microscope at UT Southwestern Live Cell Imaging Core Facility. The quantification was performed by ImageJ using at least three independent biological replicates.

### Culture and treatment of BMDM

BMDMs were harvested from eight-week-old male *Parp1* KO^LysM^ or control mice as previously described [10]. Briefly, femurs and tibias from the hind legs of the mice were used for the bone marrow extraction. Prior to bone marrow collection, the mice were euthanized using CO_2_. The bones were removed, cleaned to remove skin, muscle and cartilage, and flushed with PBS under sterile conditions to collect the bone marrow. The bone marrow suspension was filtered through a 70 μm cell strainer, and centrifuged for 10 min at 500 g at 4 °C. The cell pellets were resuspended in RBC lysis buffer to removes red blood cells. After centrifugation, the bone marrow-derived cells were resuspended in RPMI-1640 medium (Sigma-Aldrich, R8758) containing 10% fetal bovine serum (Sigma-Aldrich, F8067) and 20 ng/mL M-CSF (GenScript, Z02930). The cells were incubated for 6 days to allow the differentiation for the precursors into macrophages. On day 7, the macrophages were used for the following experiments. BMDMs from *Parp1* KO^LysM^ or control mice were treated with 500 ng/ml LPS (Sigma-Aldrich, L2630), respectively. The cells were harvested at 0, 1, 6, and 12 hour upon LPS treatment.

***Generation of conditioned medium for preadipocyte culture.*** BMDMs from *Parp1* KO^LysM^ or control mice were treated with 100 ng/ml LPS for 4 hours, then washed with PBS and replaced with fresh culture medium. After 12 hours of culture, the conditioned medium was collected from *Parp1* KO^LysM^ or control BMDMs, respectively, and used for preadipocyte culture and differentiation.

### Culture and differentiation of primary preadipocyte

Culture and differntiation of primary preadipocytes from stromal vascular fraction (SVF) of white adipose tissue were described previously [17]. Briefly, the dissected white adipose tissue from 8-week-old male wild-type C57BL/6 mice were washed, minced, and then digested for 1 hour at 37 C in PBS containing 0.1% (w/v) collagenase type I (Sigma-Aldrich, SCR103). Digested tissue was then filtered through a 100 μm cell strainer, and centrifuged at 1500*g* for 5 min to pellet the SVF cells. The SVF cells were cultured in DMEM (Cellgro, 10-017-CM) supplemented with 10% fetal bovine serum (Sigma-Aldrich, F8067) and 1% penicillin/streptomycin. At confluence, cells were induced differentiation by the addition of an adipogenic cocktail containing 1 μM dexamethasone (Sigma-Aldrich, D4902), 0.25 mM IBMX (3-isobutyl-1-methylxanthine; Sigma-Aldrich, 410957), 10 μg/mL insulin (Biosharp, BS901), and 125 nM indomethacin (Sigma-Aldrich, I7378). Two days after induction, cells were maintained in culture medium containing 10 μg/mL insulin (5 μg/ml) and 125 nM indomethacin until harvest. For coculture with the conditioned medium during differention, the SVF cells were cultured with 1:1 ratio of differention medium and the conditioned medium collected from LPS-treated BMDMs described above, throughout the period of differentiaon.

### Oil red O staining

Primary preadipocytes were differentiated as described above. After 8 days of differentiation, the cells were rinsed twice with PBS and fixed with 4% paraformaldehyde. The fixed cells were incubated with 60% isopropanol for 5 minutes and then stained with 0.3% oil red-O solution (Sigma-Aldrich, O1391) for 5 minutes.

### Flow cytometry

*Parp1* KO^LysM^ and control mice were fed with HFD for 12 weeks. SVF was isolated from visceral white adipose tissue of the mice as described above. Immune cells from SVF were characterized by flow cytometry using the following fluorochrome-conjugated antibodies targeting specific surface proteins: CD45-APC/Cy7 (BD Biosciences, 557659; RRID:AB_396774), CD11b-FITC (BD Bioscience, 557396; RRID:AB_396679), and F4/80-BV421 (BD Bioscience, 565411; RRID:AB_2734779). Cells were analyzed using a FORTESSA X20 flow cytometer (Becton Dickinson, New Jersey). The results were acquired using the FACS Diva software (BD Biosciences). Macrophages (CD45^+^/CD11b^+^/F4/80^+^) were isolated using a BD Influx cell sorter (BD Biosciences).

### Preparation of whole cell lysates

Cells were collected, washed with ice-cold PBS, and lysed in RIPA buffer (50 mM Tris-HCl pH 7.5, 450 mM NaCl, 1% NP-40, 0.1% SDS, 0.5% deoxycholic and cocktail protease inhibitors) for 15 minutes on ice. The cell lysates were clarified by centrifugation at 14000g for 15 min at 4°C. Protein concentrations were measured using the Bio-Rad Protein Assay Dye Reagent and volumes of lysates containing equal total amounts of protein were mixed with 4x SDS loading dye with Bromophenol blue. The lysates were then incubated for 5 min at 100°C.

### Western blotting

Equal amounts of whole cell lysates were prepared and run on 10% polyacrylamide-SDS gels. The gels were then transferred to a polyvinylidene fluoride membranes and blocked with 5% nonfat milk in Tris-Buffered Saline with 0.1% Tween 20 (TBST) for 1 hr at room temperature. Primary antibodies were diluted in TBST and incubated at 4°C overnight. The membranes were washed multiple times in TBST and then incubated with horseradish peroxidase (HRP)-conjugated secondary antibody diluted in 5% nonfat milk in TBST for 1 hr at room temperature. Signals were captured using a luminol-based enhanced chemiluminescence HRP substrate (SuperSignal™ West Pico, Thermo Scientific) and a ChemiDoc imaging system (Bio-Rad).

### Single cell RNA-sequencing (scRNA-seq)

***Cell preparation***. *Parp1* KO^LysM^ and control mice were fed with HFD for 12 weeks. Visceral white adipose tissue were collected, washed with ice-cold PBS, and dissociated using the SeekMate Tissue Dissociation Reagent Kit A Pro (SeekGene, K01801301). CD45^+^ immune cells were sorted by flow cytometry as described above. Cell counting and viability was analyzed using a fluorescence cell analyzer (Countstar Rigel S2) with AO/PI reagent. Then cells were washed twice with RPMI-1640 medium (Gibco, 11875119), and resuspended at 1×10^6^ cells / mL in RPMI-1640 medium supplemented with 2% FBS (Gibco, 10100147C).

***Library construction and sequencing.*** Single-cell RNA-Seq libraries were prepared using SeekOne^®^ Digital Droplet Single Cell 3’ library preparation kit (SeekGene, Catalog No. K00202). Briefly, appropriate number of cells were mixed with reverse transcription reagent and then added to the sample well in SeekOne^®^ chip S3. Subsequently, barcoded hydrogel beads (BHBs) and partitioning oil were dispensed into corresponding wells in chip S3. After emulsion droplet generation, reverse transcription were performed at 42℃ for 90 minutes and inactivated at 85℃ for 5 minutes. Next, cDNA was purified from broken droplet and amplified in PCR reaction. The amplified cDNA product was then cleaned, fragmented, end repaired, A-tailed and ligated to sequencing adaptor. The indexed PCR was then performed to generate sequencing libraries. After purification with VAHTS DNA clean beads (Vazyme, N411-01), the scRNA-seq libraries were subjected to QC analyses (i.e. the final library yield, and the size distribution of final library DNA fragments) and sequenced using an Illumina NovaSeq 6000.

***Data analysis.*** Data analysis for scRNA-seq was performed by the Green Center Computational Core Facility at UT Southwestern Medical Center. scRNA-seq libraries were demultiplexed and mapped to the mouse reference genome (mm10) using the Cellranger software (ver 3.1.0, 10× Genomics) [1]. (1) Normalization and Log-transformation: The total UMI counts per cell were normalized to the median library size across all cells using total-count scaling to 10,000 counts per cell in Scanpy v1.9.6 [2]. The normalized matrix was then log-transformed using the natural logarithm (log1p) with a pseudocount of 1. (2) Batch correction: Batch effects between *Parp1* KO*^LysM^* and control conditions were mitigated by concatenating datasets and integrating the condition as a batch label during joint preprocessing. (3) Principal component analysis (PCA): PCA was performed on the log-transformed matrix using Scanpy’s implementation built on scikit-learn v1.4 [3]. Gene expression was centered and scaled so that each gene had mean 0 and variance 1 across all cells prior to dimensionality reduction. The top principal components were used for downstream neighborhood graph construction and visualization. (4) t-SNE visualization: Two-dimensional non-linear embedding of cells was computed using the Barnes-Hut implementation of the t-SNE algorithm [4] via Scanpy, with perplexity set to 30 and default settings. The input to t-SNE was the top principal components computed earlier. (5) Cell type assignment using marker gene expression: Cluster-level cell type annotations were inferred based on curated immune and stromal marker genes. For each Leiden cluster, the mean expression of marker genes was computed per cell type. The cluster was annotated with the cell type whose marker set had the highest average expression. (6) Differential gene expression analysis: Differential expression between *Parp1* KO*^LysM^* and control cell populations was performed using the Wilcoxon rank-sum test implemented in sc.tl.rank_genes_groups. Genes with FC>1.5 and a p-value < 0.05 were considered significantly regulated. (7) Weighed *Parp1* gene expression analysis: A global scaling normalization approach was applied using ‘sc.pp.normalize_total’to scale each cell’s total transcript count to 10,000. Following this, a log-transformation with a pseudocount of 1 was applied using ‘sc.pp.log1p’ to reduce variance and normalize the expression distribution. Expression values for the gene of interest (*Parp1*) were extracted post-normalization. Each cell was associated with a replicate identifier from the metadata (obs[“replicate”]). Expression values were then summed across all cells within each replicate, providing a total expression measure per replicate.

***Gene ontology analysis.*** Gene Ontology analysis was conducted to identify enriched biological processes, KEGG pathways, and cellular components amongst experimental conditions, by using DAVID (Database for Annotation, Visualization, and Integrated Discovery) tool [69].

***CellChat-based intercellular communication analysis.*** Cell communication analysis was performed using CellChat v1.6 [39]. The processed annData object was converted into two matrices: for downstream analysis with CellChat v1.6: (1) Expression Matrix: A genes × cells matrix with gene names as row indices and cell barcodes as columns. Since the matrix was sparse, it was converted to dense format using scipy. (2) Metadata Table: A cell-wise annotation table containing the assigned cell type (Cell_Type) and experimental condition (Condition) for each cell. These matrices were saved as .csv files for downstream signaling inference using CellChat in R. The mouse ligand-receptor interaction database CellChatDB.mouse was loaded and assigned to the @DB slot of the CellChat object. The data were subset to include only expressed genes and valid ligand-receptor interactions using subsetData. Overexpressed genes and interactions were identified using ‘identifyOverExpressedGenes’ and ‘identifyOverExpressedInteractions’. Protein-protein interaction network data (PPI.mouse) were projected onto the expression dataset using ‘projectData’. Communication probabilities were inferred using ‘computeCommunProb’ with raw.use = TRUE and adjusted for population sizes. Low-confidence interactions were filtered using filterCommunication (min.cells=3), and pathway-level signaling probabilities were estimated using ‘computeCommunProbPathway’. Final network aggregation was performed with a signaling strength threshold of 0.001 using ‘aggregateNet’. Multiple visualization strategies were used to summarize the inferred communication networks: (1) Bubble plots were generated using ‘netVisual_bubble’ to show pathway-specific signaling with selected source and target cell types. (2) Chord diagrams were generated with ‘netVisual_chord_gene’ for gene-level representation of ligand-receptor interactions within individual signaling pathways.

### Quantification and statistical analyses

Untargeted metabolomics was performed with four independent biological samples, and scRNA-seq was performed with three independent biological samples. Statistical analyses for the metabolomic and genomic experiments were performed using standard statistical tests as described above. All gene-specific qPCR-based experiments were performed a minimum of three times with independent biological samples. All Western blotting experiments were performed a minimum of three times with independent biological samples. Statistical analyses were performed using GraphPad Prism 10. All tests and p values are provided in the corresponding figures or figure legends.

### Data availability

All raw data and information generated for this study are available upon request from the corresponding authors (DH and WLK) and the public data repositories noted below. The types of data and information available are as follows:

1) The scRNA-seq dataset generated for this study can be accessed from the NCBI’s Gene Expression Omnibus (GEO) repository (http://www.ncbi.nlm.nih.gov/geo) using the following accession numbers: GSE300374.
2) The raw data from the metabolomics analyses are held by Metware Biotechnology, Inc., Wuhan, China. Processed data, including metabolite lists and abundances from specific analyses can be obtained by contacting the corresponding authors (DH and WLK).
3) Software, scripts and other information about the analyses can be obtained by contacting the corresponding authors (DH and WLK).

## Acknowledgements

The authors would like to thank the members of the Huang and Kraus labs for continued input and feedback on this project.

## Author Contributions

D.H. and W.L.K. conceived and developed this project, designed the experiments, and oversaw their execution. J.Z. preformed most of the wet lab experiments and analyzed the data. R.G. assisted with the design of the mouse studies and performed some of the experiments in mice. T.N. performed the scRNA-seq data analysis. K.H provided support for some of the mouse studies. D.H. and W.L.K. prepared, edited, and finalized the figures and text, with input from other authors. W.L.K. and D.H. secured funding to support this project, and provided intellectual support and overall leadership for the work.

## Supplementary data

Supplementary data are included with this article. They contain Supplementary Figures S1 through S4.

## Supplementary Figure Legends

**Figure S1.**
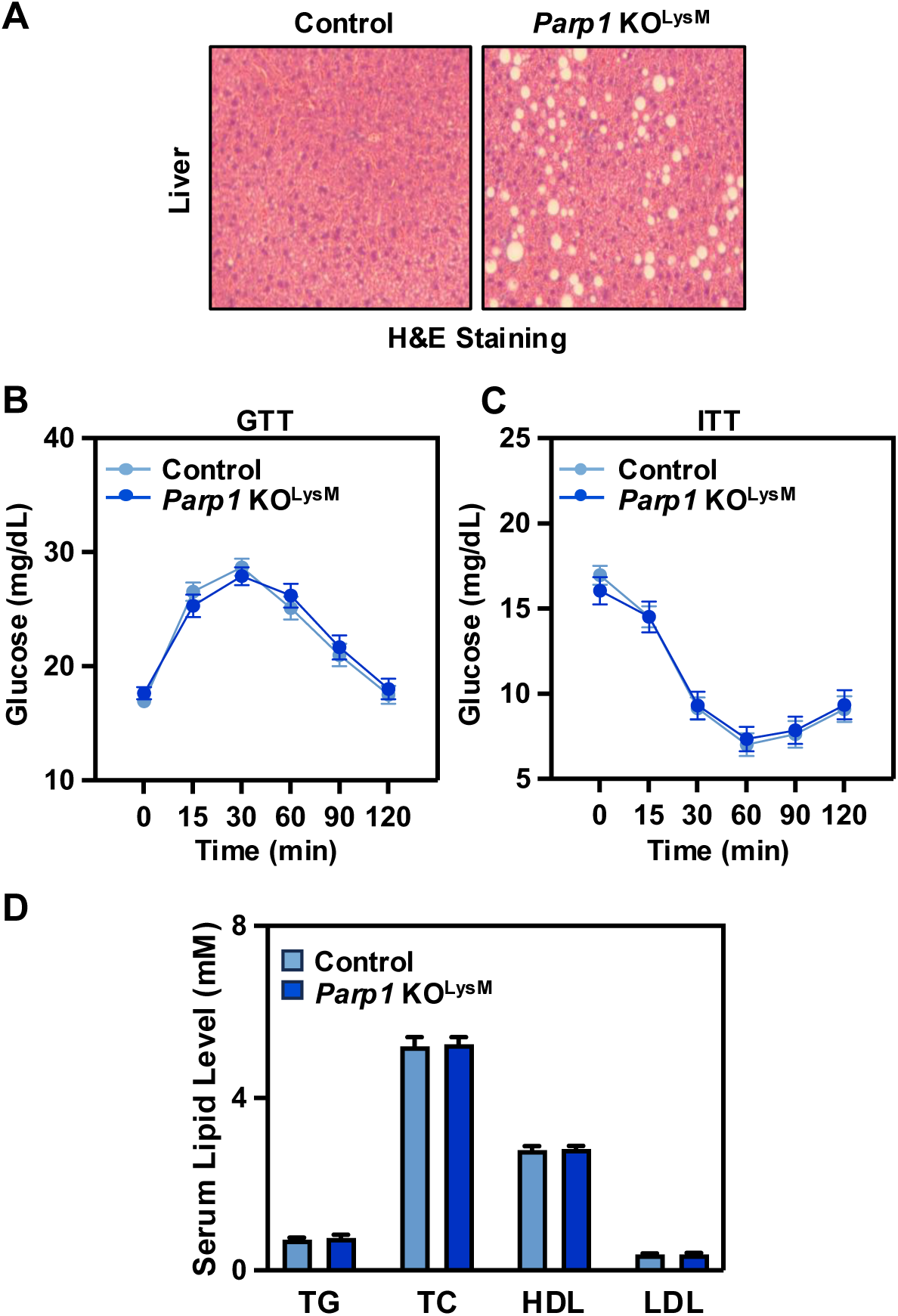
Effects of macrophage-specific depletion of PARP1 on obesity-associated metabolic outcomes. (**A**) Representative images of H&E staining showing lipid accumulation in liver sections from control and *Parp1* KO^LysM^ mice after 8 weeks of HFD feeding. **(B)** Results of glucose tolerance test (GTT) for control and *Parp1* KO^LysM^ mice after 12 weeks of HFD feeding. Each point represents the mean ± SEM (control, n = 25 for each group; *Parp1* KO^LysM^, n = 28 for each group). No significant differences analyzed by unpaired t-test. **(C)** Results of insulin tolerance test (ITT) for control and *Parp1* KO^LysM^ mice after 12 weeks of HFD feeding. Each point represents the mean ± SEM (control, n = 25 for each group; *Parp1* KO^LysM^, n = 28 for each group). No significant differences analyzed by unpaired t-test. **(D)** Serum lipid analyses for control and *Parp1* KO^LysM^ mice after 12 weeks of HFD feeding. Each bar represents the mean + SEM (control, n = 25 for each group; *Parp1* KO^LysM^, n = 28 for each group). No significant differences analyzed by unpaired t-test.

**Figure S2.**
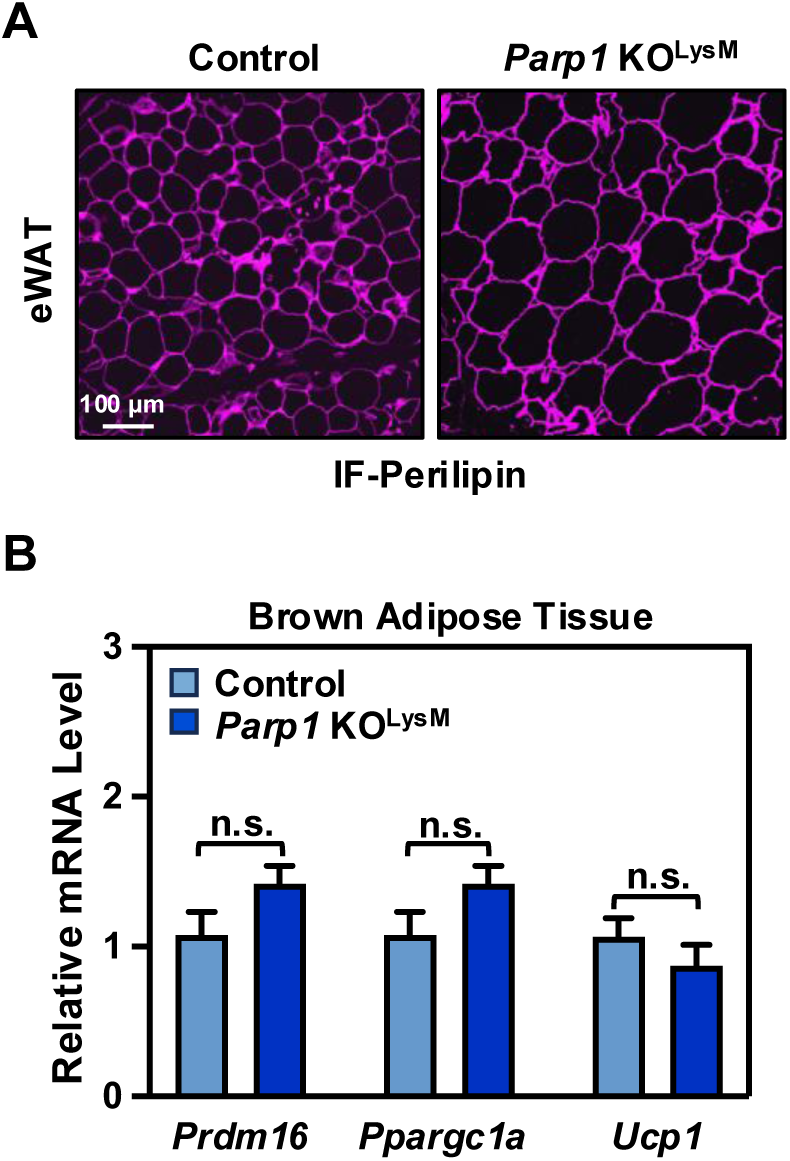
Effects of macrophage-specific depletion of PARP1 on adipose tissues during diet-induced obesity. **(A)** Immunofluorescent staining of perilipin in eWAT from control and *Parp1* KO^LysM^ mice after 12 weeks of HFD feeding. **(B)** Expression of thermogenic genes *Prdm16, Ppargc1a,* and *Ucp1* in brown adipose tissue, as determined by RT-qPCR. Each bar represents the mean + SEM (n = 10 for each group). No significant differences analyzed by unpaired t-test.

**Figure S3.**
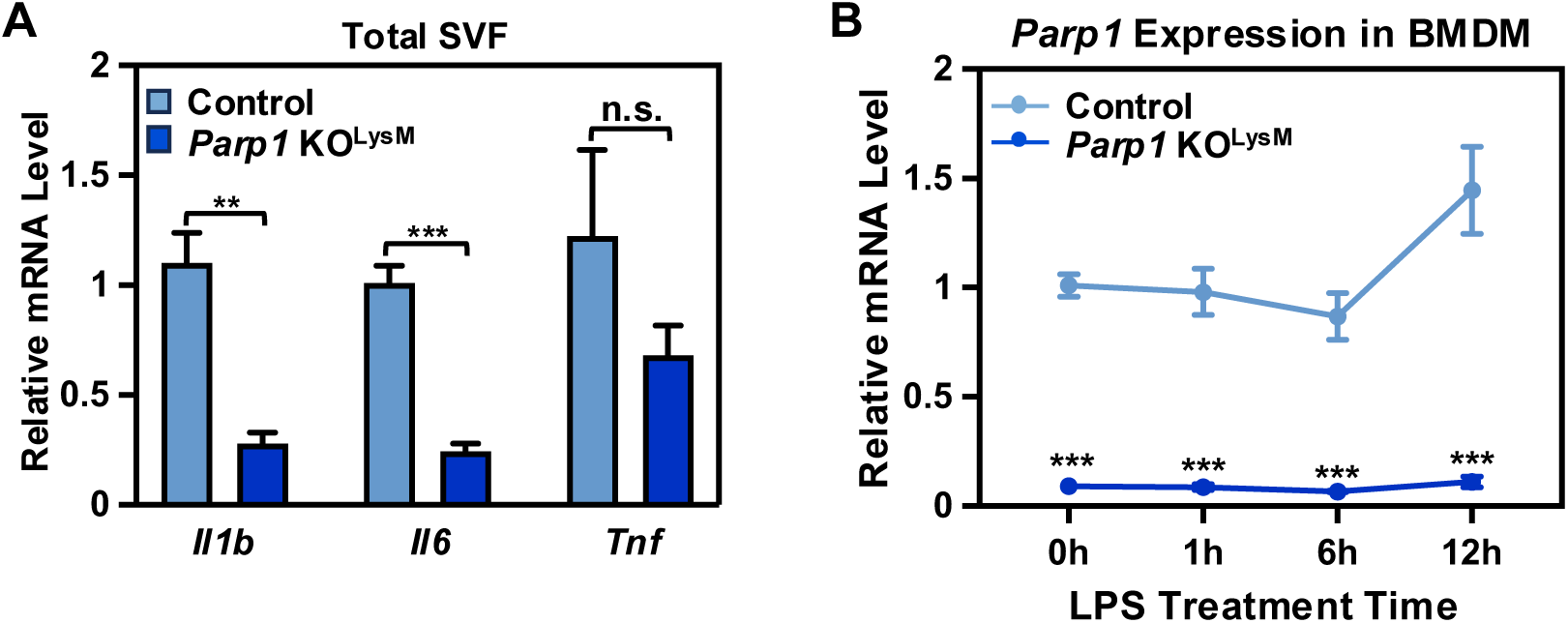
Depletion of PARP1 in macrophages alleviates inflammatory responses in WAT. **(A)** Expression of proinflammatory genes *Il1b, Il6,* and *Tnf* in SVF isolated from eWAT of control and *Parp1* KO^LysM^ mice, as determined by RT-qPCR. Each bar represents the mean + SEM (n = 4 for each group). Asterisks indicate significant differences from the control; two tailed, unpaired t-test; **, p < 0.05; and ***, p < 0.001; n.s., no significant difference. **(B)** Expression of *Parp1* gene in BMDMs isolated from control and *Parp1* KO^LysM^ mice, upon treatment with LPS, as determined by RT-qPCR.. Each point represents the mean ± SEM (n = 9 for each group). Asterisks indicate significant differences from the control at individual time points; two tailed, unpaired t-test; ***, p < 0.001.

**Figure S4.**
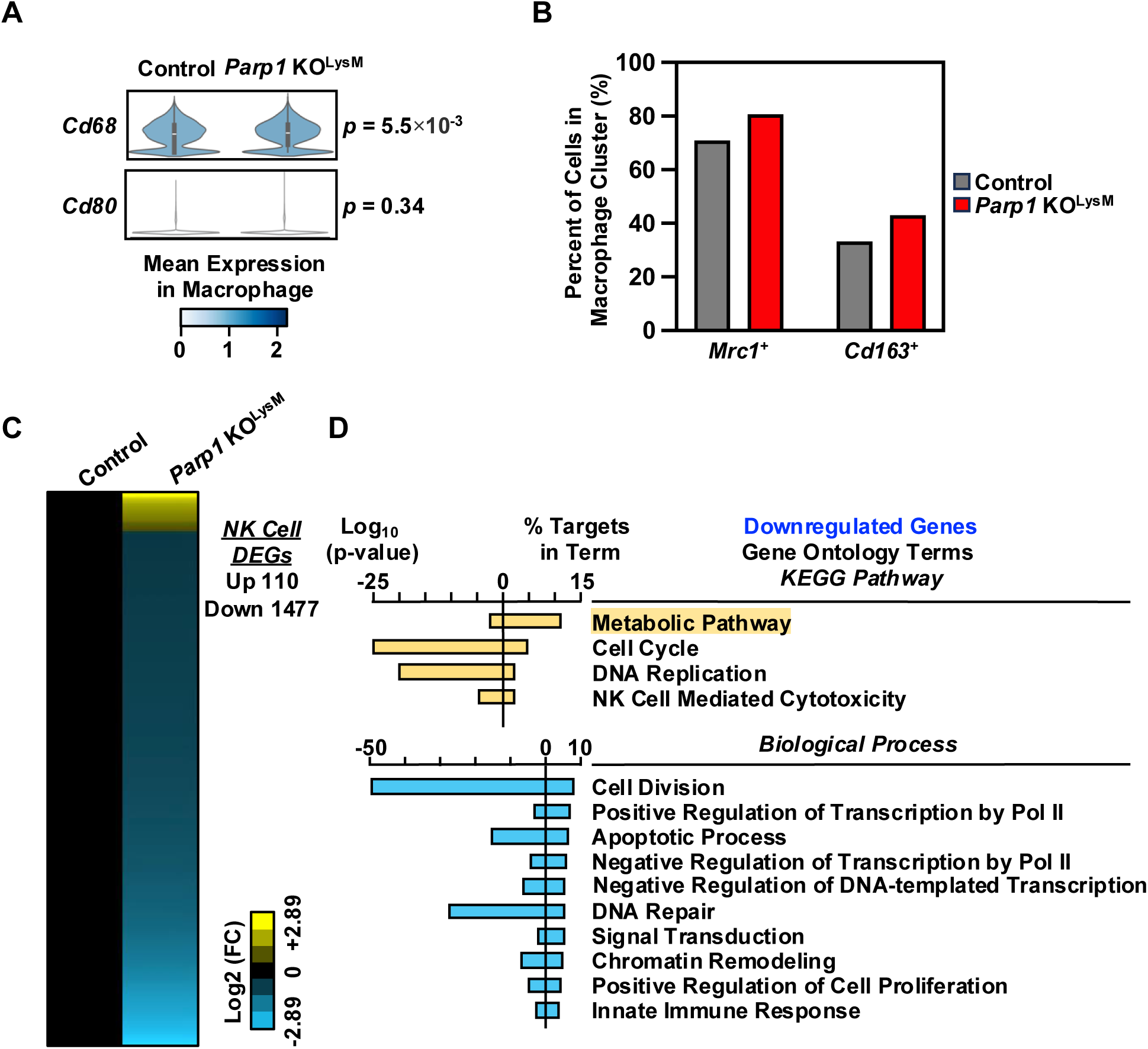
Effects of macrophage-specific depletion of PARP1 on the regulation of macrophage phenotypes and NK cell gene expression. (**A**) Violin plots showing expression level of M1-type marker genes *Cd68* and *Cd80* in macrophage cluster from scRNA-seq between control and *Parp1* KO^LysM^ groups, analyzed by Wilcoxon Signed-Rank test (p value as indicated). **(B)** Percent of cells expressing *Mrc1* or *Cd163* within macrophage cluster in control versus *Parp1* KO^LysM^ group, respectively. **(C)** Heatmaps showing differentially expressed genes in NK cell cluster between control and *Parp1* KO^LysM^ groups from scRNA-seq. **(D)** Gene Ontology terms for DEGs in NK cell cluster shown in **(C)**.

